# Tumor Cell Clustering Enhances Metastatic Competence by Regulating the H3K36 Histone Demethylase KDM2A

**DOI:** 10.64898/2026.01.22.701120

**Authors:** Carolyn Kravitz, Kiran D. Patel, Emily Wingrove, Dejian Zhao, Tang Tang, Sampada Chande, Minghui Zhao, Yuchen Huo, Thomas F. Westbrook, Qin Yan, Don X. Nguyen

## Abstract

Disseminated tumor cells can form clusters via cell-cell adhesion, which increases their capacity to initiate metastasis. Metastatic clusters are characterized by distinct changes in transcription, suggesting that epigenetic mechanisms underlie their unique phenotypic state. By performing functional epigenomic studies in models of non-small cell lung cancer, we identified the histone H3 lysine 36 (H3K36) demethylase KDM2A as being differentially required for the fitness of metastatic cell clusters. This contextual dependency on KDM2A is predicated by tumor cell-cell aggregation, which specifically induces KDM2A binding to CpG island enriched promoters. At these defined genomic loci, KDM2A maintains H3K36 monomethylation, which preferentially correlates with transcriptional activation. KDM2A directly targets oxidative phosphorylation genes and KDM2A activity is required for optimal mitochondrial respiration and apical cell junction integrity in cell clusters. Consequently, suppressing KDM2A reduces metastatic seeding and colonization in multiple organs, including in the brain. These findings reveal a chromatin regulatory mechanism by which homotypic cell communication instructs the epigenome of disseminated tumor cells to potentiate their metastatic competence.

## Introduction

Disseminated tumor cells (DTCs) captured from the bloodstream of humans appear as single cells or microemboli (also referred to as “clusters”) across a wide variety of epithelial cancers ^1^. Clusters can form via homotypic tumor cell-tumor cell interactions during collective cell invasion or *de-novo* aggregation in circulation ^2^. In preclinical models, the metastatic capacity of clusters is increased relative to single CTCs or dissociated cancer cells ^3, 4^, in part because aggregation protects DTCs from intravascular oxidative stress and immune cell killing ^5^. Homotypic clustering of cancer cells is mediated by apical cell junctions (e.g. adherens junctions, tight junctions, or desmosomes) ^3, 6, 7^ as well as other cell adhesion molecules ^8, 9^. Importantly, the detection of clusters in the plasma of patients with cancer correlates with poor outcome ^10–14^. Furthermore, in patients with late-stage disease, bursts in CTC clusters may lead to macrovascular invasion and cancer associated lethality ^15^. Accordingly, clinical trials are underway to test if drugs that disrupt cell-cell junctions can be safely used to reduce DTC burden in humans ^16^.

In primary tumors, certain low frequency somatic mutations correlate with the expansion of malignant sub-clones and their inferred capacity to seed secondary metastases ^17^. Alternatively, the metastatic proclivity of cancer cells is also associated with transcriptional ^18^ and epigenetic alterations ^19–21^ that occur independently of genetic mutations. Examples include modifications in DNA and histone tails, which are regulated or interpreted by writer, eraser, or reader protein complexes ^22^. During metastasis, epigenetic changes reversibly alter transcriptional outputs to engage adaptive mechanisms by which malignant cells can disseminate, persist as dormant lesions, evade the immune system, or colonize distant organs ^23^. Chromatin remodeling in particular is a major determinant of tumor cell plasticity, which is defined as the ability of cancer cells to transition between phenotypic states in response to therapy or cues from the tumor microenvironment ^24^. Deletion of chromatin modifying proteins causes specific transcriptional readouts, likely due to their recruitment to particular genomic loci in a manner that is cell state dependent. Understanding the contextual requirement for chromatin modifiers could reveal novel targeting strategies to inhibit tumor cell plasticity and metastasis.

Dynamic changes in transcription ^25^ and DNA hypomethylation ^7^ have been linked to the formation of metastatic cell clusters, suggesting that cancer cell aggregation induces distinct epigenetic and cellular states. During tissue morphogenesis, cell adhesion itself functions as an instructive molecular cue ^26, 27^ and epithelial cell junction formation can trigger DNA methylation ^28^. Whether chromatin regulatory mechanisms are induced in response to the aggregation of DTCs and their impact on metastatic cell phenotype(s) is unknown. Herein we performed functional epigenomic studies to identify epigenetic dependencies of tumor cell clusters and elucidated their biological requirements during cancer metastasis, including in non-small cell lung cancer (NSCLC).

## Results

### Identifying epigenetic modulators of NSCLC cell fitness in metastatic clusters

The biological significance of DTC clusters has been documented in several cancer models such as melanoma ^29^ and breast cancer ^3^. By contrast, the role of tumor cell clustering in NSCLC, the most frequently diagnosed type of thoracic malignancy, remains poorly understood. Notably, peri-operative detection of DTC clusters in patients with NSCLC correlates with a decrease in recurrence-free survival ^30^. In mouse models, we and others have found that NSCLC cells can extravasate and then expand as perivascular clusters in multiple organs including the brain ^31, 32^, which is a major site of metastasis in patients with advanced stage lung cancer ^33^. The H2030-BrM3 cells are a well characterized metastatic sub-population of the human H2030 parental cell line which is prototypical of NSCLC driven by *KRAS* and *p53* mutations ^34^. When cultured under low Matrigel supplemented suspension conditions which induce tumor cell clustering in other cancer models ^25^, H2030-BrM3 cells also form aggregates consisting of 2-20 viable cells within 24 hours (**Fig. 1A)**. To test the *in vivo* metastatic capacity of NSCLC tumor cell clusters, we allowed H2030-BrM3 cells to aggregate *in vitro* and then injected cell clusters into the arterial circulation of immunocompromised mice. In parallel, a group of mice were injected with an equivalent number of dissociated cancer cells (**Fig. 1B**). Metastatic progression in multiple organs was enhanced in animals injected with H2030-BrM3 clusters as compared to mice injected with dissociated cells (**Fig. 1C**). Furthermore, tumor cell clusters had a significant increase in their capacity to seed the brain (**Fig. 1D-E**). We also performed bulk RNA-sequencing comparing H2030-BrM3 clusters to H2030-BrM3 cells grown under sub-confluent adherent conditions (referred to as control). As expected, H2030-BrM3 aggregations *in vitro* caused specific changes in gene expression over 5 days (**Extended Data Fig. 1A**).

**Figure 1:**
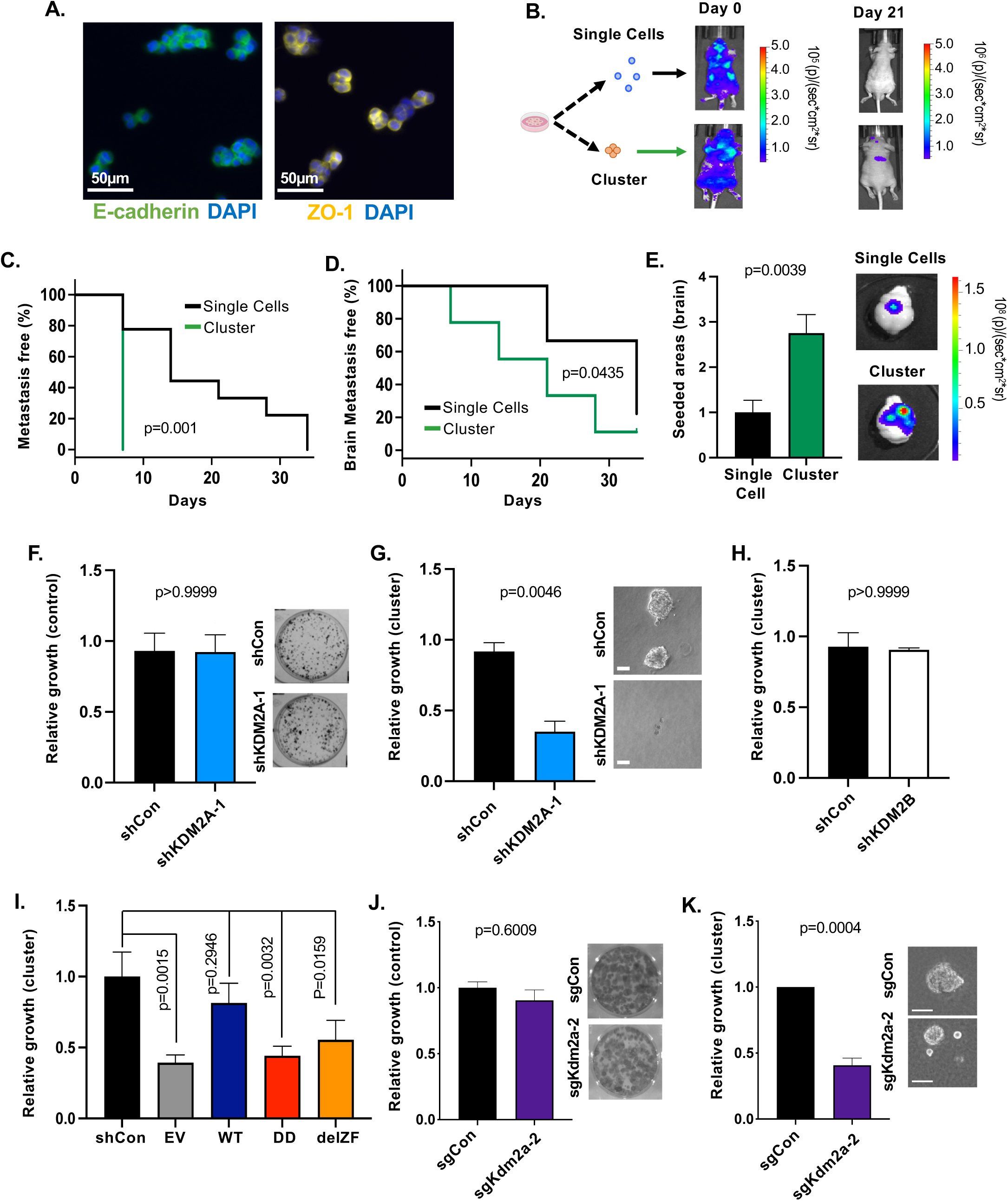
The fitness of metastatic cell clusters requires KDM2A. **(A)** Immunofluorescent staining (IF) of DAPI (blue), E-cadherin (green; left), or ZO-1 (yellow; right) in H2030-BrM3 cells 24 hours after seeding under cluster forming conditions. Scale=50µm. **(B)** Schema and representative images of animals injected with an equal number of dissociated or clustered H2030-BrM3 cells on Day 0 and Day 21 after intra-arterial injection. Metastasis was detected by bioluminescent imaging (BLI). **(C,D)** Metastasis free survival (C) and brain metastasis free survival (D) of animals injected with H2030-BrM3 cells as dissociated cells or clusters from (B). P-value by Mantel-Cox test. **(E)** Quantification (left) and representative images (right) of metastatic seeding in the brain of animals from (D). Data was quantified as distinct foci per animal at endpoint. P-value was calculated by Welch’s t-test. **(F)** Quantification (left) and representative images (right) of clonogenic growth of H2030-BrM3 cells expressing the indicated doxycycline (Dox) inducible shRNAs. Plotted are values from Dox treated samples relative to untreated shCon samples after 10 days. P-value by Mann-Whitney. N=3 biological replicates. **(G)** H2030-BrM3 cells expressing luciferase and the indicated shRNAs were cultured under cluster forming conditions. Relative cell growth was measured based on luciferase activity after 12 days and plotted for Dox treated samples relative to untreated shCon samples. P-value by Welch’s t-test. N=3 biological replicates. Representative images of clusters at endpoint are shown (right). Scale=100μm. **(H)** Relative growth of H2030-BrM3 tumor cell clusters expressing shRNAs against *KDM2B* was measured as in G. P-value by Mann-Whitney. N=2 biological replicates. **(I)** Relative growth of cell clusters co-expressing shKDM2A-1 with empty vector (EV), 3X Flag tagged full length wild type *KDM2A* (WT), 3X Flag *KDM2A (*DD), or 3X Flag *KDM2A* (delZF) was measured as in G. P-values were calculated by Fisher’s LSD test following one-way ANOVA. All adjusted p-values ≤ 0.05 by Dunnet’s multiple testing comparison. n=7 biological replicates. **(J)** Quantification (left) and representative images (right) of clonogenic growth of LU-KP cells expressing Cas9 and the indicated sgRNAs after 7 days. Data was measured as in F. P-value by one-way ANOVA with Dunnett’s multiple testing correction. N=3 biological replicates. **(K)** Growth under cluster forming conditions of LU-KP cells with the indicated sgRNAs was quantified as in G. N=5 biological replicates. Representative images of clusters at endpoint are shown (right). Scale=50μm. P-value by one-sided t-test against mean of shCon samples. Bar graphs show mean +/− standard error of the mean (SEM) unless otherwise noted.

Having determined that the aggregation of H2030-BrM3 cells increases their metastatic potential and induces distinct transcriptomic features, we performed a functional screen using this model to identify epigenetic dependencies of NSCLC clusters. We employed an inducible, barcoded short hairpin RNA (shRNA) library that was assembled to target a subset of epigenetic modulating proteins (n=89) whose function can be probed using small-molecules and/or whose expression in human cancers corelates with poor clinical outcome ^35^. The screen was then implemented using pooled cell populations, comparing barcode abundance in cell clusters to cells grown under control conditions *in vitro* after 12 days. As expected, several targets, including a control shRNA against *KRAS*, were reduced across conditions, indicating that they target essential genes (**Extended Data Fig. 1B and Supplementary Table 1**). By contrast, we also identified targets that were preferentially reduced in metastatic clusters including shRNAs against *BRD9* and *KDM2A* (cluster specific targets; **Extended Data Fig. 1B, orange dots**).

### The lysine-specific demethylase KDM2A enhances metastatic competence

Of the cluster specific targets, we were able to validate the effects of *KDM2A* knockdown. Specifically, as compared to cells expressing a non-targeting shRNA (shCon), *KDM2A* knockdown using two independent shRNAs (shKDM2A-1 and -2) (**Extended Data Fig. 1C)** did not have any effects on cell viability under control conditions (**Fig. 1F and Extended Data Fig. 1F)** but rather inhibited the outgrowth of clusters over a period of 10-12 days **(Fig. 1G and Extended Data Fig. 1E).** KDM2A is a jumonji C (JmjC) domain containing protein that demethylates the dimethyl residue of histone H3 at lysine 36 (H3K36) to generate mono-methylated H3K36 ^36^. Knockdown of *KDM2B*, the only other member of the KDM2 family, had no effect on cluster outgrowth **(Extended Data Fig. 1F and Fig. 1H),** suggesting a preferential role for KDM2A in this context. In addition to the JmjC domain, KDM2A encodes a CXXC domain, which can modulate transcription independently of histone demethylation ^37–39^. As such, we ectopically re-expressed FLAG tagged shRNA resistant cDNAs encoding for full length wild type *KDM2A* (wt), a demethylase defective mutant of *KDM2A* (DD), or a CXXC deletion mutant of *KDM2A* (delZF) into cells with endogenous *KDM2A* knockdown. Whereas expression of KDM2A(wt) rescued the outgrowth of clusters, the DD and delZF mutants failed to do so completely (**Extended Data Fig. 1G and Fig. 1I**). In addition, treatment with the KDM2/7 inhibitor (MCE cat. HY-108706), which inhibits KDM2A demethylase activity at 0.1-1µM ^40^, also decreased cluster outgrowth (**Extended Data Fig. 1H**). Next, we reduced Kdm2a via CRISPR-Cas9 gene editing in a murine cancer cell line (LU-KP) that we derived from a genetically engineered mouse model of NSCLC driven by *KRAS* and *Tp53* ^41^ (**Extended Data Fig. 1I)**. *Kdm2a* knockdown also preferentially decreased cluster outgrowth in this independent model of NSCLC **(Fig. 1J,K and Extended Data Fig. 1J,K).**

It has been previously suggested that NSCLC tumorigenesis can be driven by *KDM2A* amplifications ^42^. However, the regulatory context of KDM2A activity and its requirement during metastatic progression to relevant distant sites are not known. Hence, we tested the *in vivo* consequence(s) of KDM2A knockdown in syngeneic and xenograft models of metastasis. First, we injected LU-KP cells into the flank of C57BL/6 mice and found that *Kdm2a* knockdown had no effect on subcutaneous tumor growth (**Fig. 2A)** but decreased the number of spontaneously formed lung nodules in the same animals (**Fig. 2B,C and Extended Data Fig. 2A**). After injecting H2030-BrM3 cells into the trachea of athymic mice, *KDM2A* knockdown had modest effects on lung orthotopic tumorigenesis at late time points (**Fig. 2D**). However, animals injected with shKDM2A cells into their lungs had a reduced frequency of spontaneous brain and/or liver metastasis at necropsy (**Fig. 2E,F**). This reduction in distant metastasis was independent of orthotopic lung tumor burden (Spearman correlation p=0.4138; **Extended Data Fig. 2B**). KDM2A is thus required for the viability of metastatic cell clusters and enhances their metastatic competence *in vivo* across murine and human models of NSCLC.

**Figure 2:**
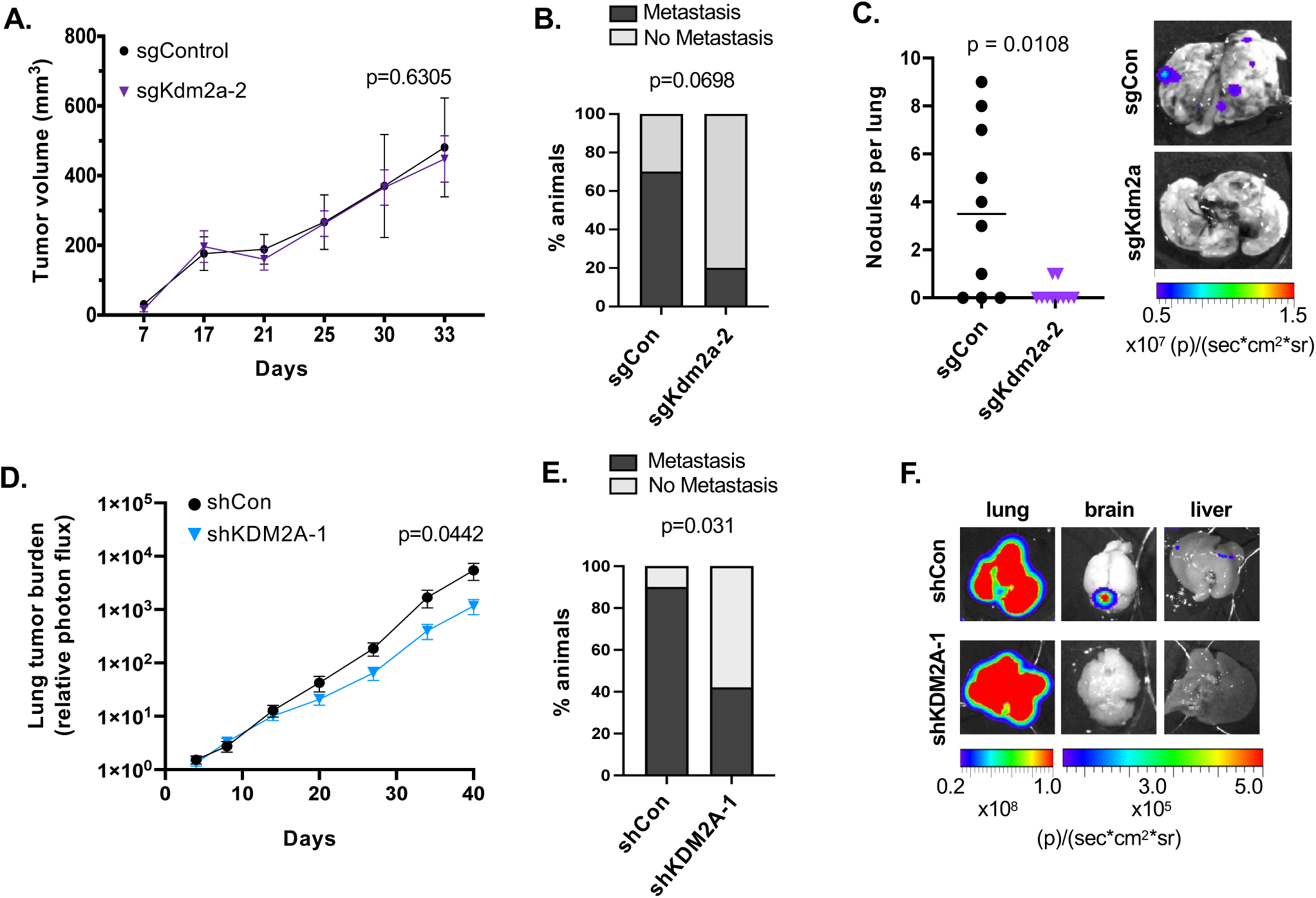
KDM2A enhances NSCLC metastasis *in vivo*. **(A)** sgCon or sgKdm2a-2 expressing LU-KP cells were injected into the flank of wild-type C57BL/6 mice, and subcutaneous tumor growth was measured. P-value for final time point by Mann-Whitney. **(B)** Incidence of lung metastasis in animals from A. P-value by Fishers exact test. N=10 animals per group. **(C)** Quantification (left) and representative images (right) of lung nodules detectable at necropsy in animals from A. P-value by Mann-Whitney. N=10 animals per group. **(D)** Dox inducible shCon or shKDM2A-1 expressing H2030-BrM3 cells were injected into the trachea of athymic mice and animals treated with Dox after Day 7. Orthotopic lung tumor burden was measured by BLI and normalized to Day 0. P-value was calculated by Mann-Whitney for Day 49. **(E)** Incidence of detectable metastatic lesions in the brain or liver of animals in D at necropsy (day 61-85). P-value by Fisher’s-exact test. N=10 control, N=12 shKDM2A-1. **(F)** Representative images and BLI at necropsy of brain and liver from animals in E. Bar graphs show mean +/− SEM unless otherwise noted.

### Tumor cell clustering induces KDM2A recruitment to CpG island (CGIs) promoters

The steady state levels of endogenous KDM2A or rescued FLAG-KDM2Awt protein did not change in H2030-BrM3 clusters (**Extended Data Fig. 3A,B**). Furthermore, the localization of KDM2A remained in the nucleus across conditions (**Extended Data Fig. 3C**). Consequently, we performed chromatin immunoprecipitation sequencing (ChIP-seq), leveraging H2030-BrM3 cells with rescued expression of FLAG-KDM2Awt. Under control conditions, KDM2A was bound to 977 genomic regions. By contrast, there was a >10-fold increase in KDM2A bound regions (n=11035) in cell clusters after 5 days (**Fig. 3A)**. Based on genomic annotation using HOMER, KDM2A is mainly located at intronic and intergenic regions in control cells, whereas KDM2A binding in tumor cell clusters specifically increases in exons and promoter regions (**Fig. 3B-D**). We next focused on gene linked loci and found that, while KDM2A can target a few common genes across conditions, the majority of KDM2A targets are bound *de novo* in metastatic clusters (**Fig. 3E**). These KDM2A bound genes encode for biological features (e.g. neurogenesis and metabolic processes) which have been previously linked to metastatic cells (**Fig. 3F and Supplementary Table 2**). Significantly, DNA binding motif enrichment analysis showed that the induced KDM2A bound sites are CG rich and predominantly linked to CpG islands (CGIs) (**Fig. 3G, H**), consistent with the capacity of KDM2 paralogs to directly bind to narrow regions of unmethylated DNA ^39^.

**Figure 3:**
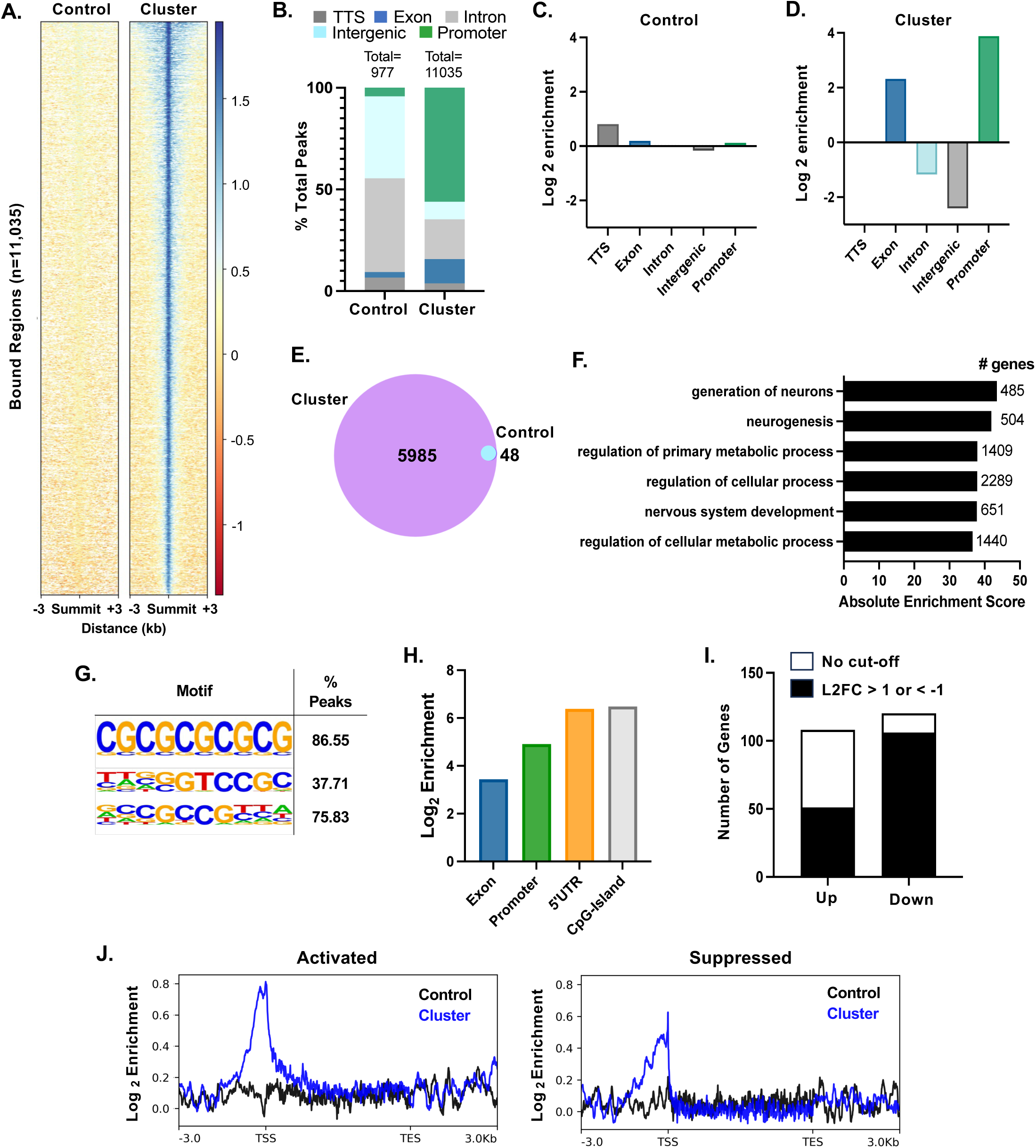
KDM2A binding to CGI enriched promoters is increased in metastatic cell clusters. **(A)** Heatmap plotting the relative peak summit binding of FLAG-KDM2A to chromatin regions in H2030-BrM3 cells under control (left) or cluster forming (right) conditions after 5 days. N=11,035 peaks, sum of all peaks across N=2 biological replicates. **(B)** Proportion of peak summits annotated by HOMER as being within promoter, intergenic, intronic, exonic, or transcription termination site (TTS) regions, in control or cluster forming conditions. Proportions represent the sum of peaks between N=2 biological replicates. N=977 control peaks, N=11035 cluster peaks. **(C)** Log2 enrichment of summit location for control and **(D)** cluster peaks shown in **(E)** Venn diagram depicting number of gene linked peaks bound by FLAG-KDM2A under control and cluster conditions from B-D. **(F)** Top Gene Ontology: Biological Pathway enrichment terms associated with KDM2A bound peaks in tumor cell clusters from (A). Absolute Enrichment Score represents –log10 of calculated enrichment. P-values <0.01 with Bonferroni correction for all. **(G)** Top three (out of 123) KDM2A binding motifs by significance were identified using HOMER. Q < 0.001 for all. No motifs reach significance in control condition. **(H)** Plotted are the Log2 enrichments of KDM2A binding peaks in tumor cell clusters and classified by indicated regulatory elements. 5’UTR = 5’ Untranscribed Region. **(I)** Number of direct KDM2A target genes that are differentially expressed between shCon and shKDM2A-1 H2030-BrM3 cells, grown under cluster forming conditions after 5 days. No direct differentially expressed genes were detected under control culture conditions. Significance determined by adjusted p-value <0.05 with Benjamini-Hochberg correction. N=2-3 biological replicates. **(J)** Profile plots of average KDM2A enrichment (defined as log2 ratio of IP:Input) at KDM2A activated and KDM2A suppressed genes in H2030-BrM3 cells under control (black) and cluster (blue) conditions. Traces are average of N=2 biological replicates.

To determine the transcriptional consequences of KDM2A activity in tumor cell clusters, we next integrated RNA sequencing analysis, comparing shKDM2A cells to shCon cells. Under control conditions, *KDM2A* knockdown had modest effects on the transcriptome of cells (n= 18 genes). Conversely, KDM2A knockdown dysregulated the expression of ∼426 genes in clusters after 5 days (**Extended Data Fig. 3D** and **Supplementary Table 3).** The genomic distribution of KDM2A binding across activated (i.e. downregulated in shKDM2A vs shCon cells) or repressed genes (i.e. upregulated in shKDM2A vs shCon cells) was similarly biased towards promoters (**Extended Data Fig. 3E**). Nevertheless, when considering KDM2A bound genes whose expression are most affected by KDM2A levels, a greater proportion of these targets are transcriptionally activated as compared to loci that may be repressed (**Fig. 3I**). Also, KDM2A binding was stronger and more centered around the transcription start sites (TSSs) of transcriptionally activated genes (**Fig. 3J**). We conclude that cell aggregation induces KDM2A binding to promoter CGIs, which correlates preferentially, although not exclusively, with transcriptional activation in metastatic cells.

### KDM2A maintains H3K36 monomethylation in active promoters

The various H3K36 methylation states are enriched in distinct regions of the genome ^43^. To ascertain if KDM2A controls H3K36 methylation and the consequences of this regulation on transcription, we performed ChIP-seq of H3K36me1-3 in H2030-BrM3 cells under control or cluster forming conditions. We focused on analyzing promoters and loci immediately preceding TSSs as these regions are preferentially bound by KDM2A. The accumulation of H3K36me1 at genic promoters was more elevated in clusters **(Fig. 4A,B)** and correlated with increasing H3K27 acetylation **(Extended Data Fig. 4A)** as well as higher gene expression **(Fig 4C).** These data indicate that, under cluster forming conditions, H3K36me1 accumulation is preferentially linked to features of transcriptionally permissive promoters. A reciprocal pattern of H3K36me2/3 was observed in the same promoter regions under control vs cluster forming conditions (**Extended Data Fig. 4B-E)**. Knockdown of KDM2A in tumor cell clusters resulted in a general reduction of H3K36me1 and increase of H3K36me2 at promoter spanning regions (**Fig. 4D,E)** as well at CpGs (**Fig. 4F,G**). These loci specific changes occurred without any differences in the overall levels (**Extended Data Fig. 4F**) or global genomic distribution (**Extended Data Fig. 4G**) of H3K36me1-3 methylation. Moreover, KDM2A mediated changes in H3K36me1/me2 were more exacerbated in KDM2A activated promoter vs. regions that are potentially repressed by KDM2A (**Fig. 4H-J, and Extended Fig. 4H**). Thus, in the same biological context, KDM2A potentiates promoter activation in a demethylase dependent manner but may constrain transcription of a smaller subset of loci independently of changes in H3K36 methylation. Altogether, these data suggest that KDM2A maintains H3K36 monomethylation in a subset of transcriptionally permissive CGI promoters.

**Figure 4:**
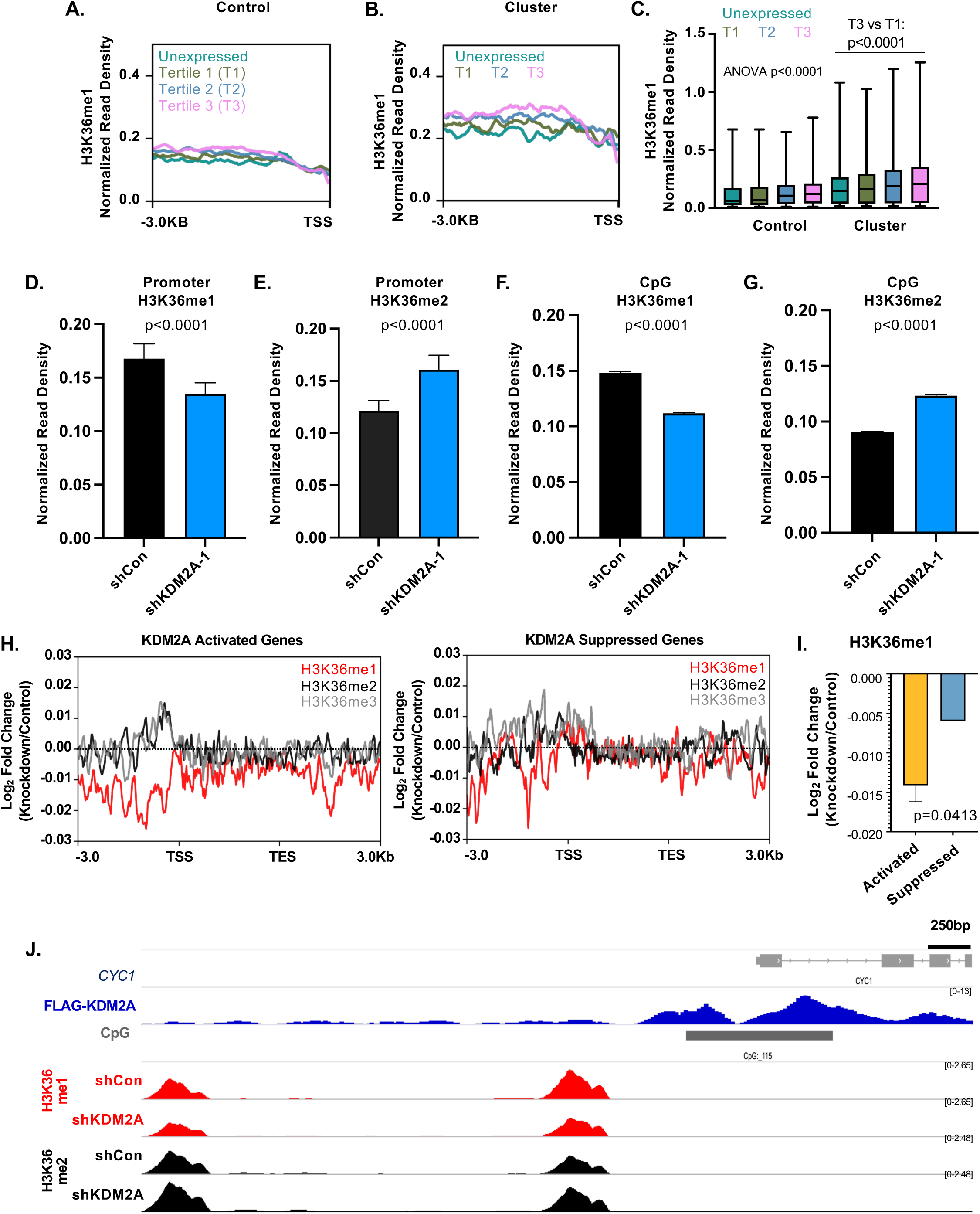
KDM2A maintains H3K36me1 at promoters and TSS of activated genes. **(A,B)** Profile plot of H3K36me1 counts per million (CPM) normalized read density (NRD) through the promoter region (−3kB to TSS) of genes in H2030-BrM3 cells, under control (A) or cluster forming (B) conditions and categorized by levels of gene expression. mRNA levels were scaled on expression in H2030-BrM3 control or cluster forming conditions taken at Day 5 ranging from unexpressed (Unex.) < Tertile 1 (T1) < Tertile 2 (T2) < Tertile 1 (T3). Number of genes per tertile are: N=3642 (Unex.), 5397 (T1), 5398 (T2), and 5399 (T3) for control and N=4163 Unex., 5219 (T1), 5235 (T2), 5219 (T3) for clusters. Traces show average of n=4 biological replicates. **(C)** Box- and-Whisker plots of NRD of traces for A and B. P-value by Kruskal-Wallis test with all comparisons p<0.0001 using Dunn’s multiple testing correction except: T0 cluster vs. T1 cluster (p=0.0059), T0 cluster vs. T3 control (p=0.0022), T0 control vs. T1 control (p=0.0039), and T2 cluster vs. T3 cluster (p=0.0007). **(D,E)** Quantification of H3K36me1 (D) or H3K36me2 (E) NRD in H2030-BrM3 clusters with the indicated shRNAs through the promoter of activated regions. N=105 loci. P-values by Wilcoxon matched-pairs test. **(F,G)** Quantification as in D,E, but in 3kB regions surrounding CpG islands genome-wide. N= 20,079 Me1 shControl, 22,200 me1 shKDM2A-1, 24,440 me2 shCon, 23,306 me2 shKdm2a-1. P-values by Wilcoxon matched-pairs test only on those loci with coverage in both conditions. **(H)** Profile plot of H3K36me1/2/3 average enrichment in KDM2A target genes that are activated (left) or suppressed (right) in H2030-BrM3 clusters. Traces are average of N=4 biological replicates collected at Day 5. **(I)** Comparative quantification of the promoter region (−3.0 Kb to TSS) for genes from both groups plotted in D. P-value by Mann-Whitney. **(J)** IGV tracks of CPM NRD for the KDM2A activated gene *CYC1* with annotation of CGI, KDM2A chromatin binding peaks, and H3K36me1 peaks in shCon and shKDM2A-1 expressing H2030-BrM3 clusters. Traces are average of n=2 for FLAG-KDM2A and N=4 for H3K36 biological replicates.

### KDM2A regulates oxidative phosphorylation and a metabolic stress program in tumor cell clusters

To identify the proximal biological consequences of KDM2A inhibition, we focused on genes that are activated by KDM2A in metastatic clusters. When comparing the transcriptome of shKDM2A clusters to shCon clusters, the most significantly depleted gene set included regulators of oxidative phosphorylation, with a high proportion of these potentially bound by KDM2A at their promoters (**Fig. 5A, Extended Data Fig. 5A and Supplementary Table 4)**. The expression of several of these genes is decreased specifically when KDM2A is inhibited in tumor cell clusters (**Fig. 5A,B**). A congruent decrease in metabolic gene expression was observed in the LU-KP cell cluster model after Kdm2a knockdown (**Fig. 5C).** The transcriptional repression of this gene set is linked to a loss of H3K36me1 and concomitant increase in H3K36me2 at their TSSs spanning regions (**Fig. 5D**). Based on these epigenetic findings, we hypothesized that a disruption of mitochondrial function may be driving the compromised cluster cell fitness following KDM2A knockdown. Hence, we subjected shCon and shKDM2A tumor cells to a mitochondrial stress test. KDM2A inhibition in cell clusters disrupted mitochondrial oxygen consumption, with significant decreases in basal respiration and maximal respiratory capacity after uncoupling with FCCP (carbonyl cyanide-p-trifluoromethoxyphenylhydrazone) (**Fig. 5E-G).** The basal respiration defects in the H2030-BrM3 clusters could be rescued by ectopic expression of KDM2Awt but not the DD and delZF mutants (**Fig. 5H**). A similar defect in respiratory capacity was observed in the LU-KP cell clusters after Kdm2a knockdown **(Extended Data Fig. 5B and Fig. 5I,J)**.

**Figure 5:**
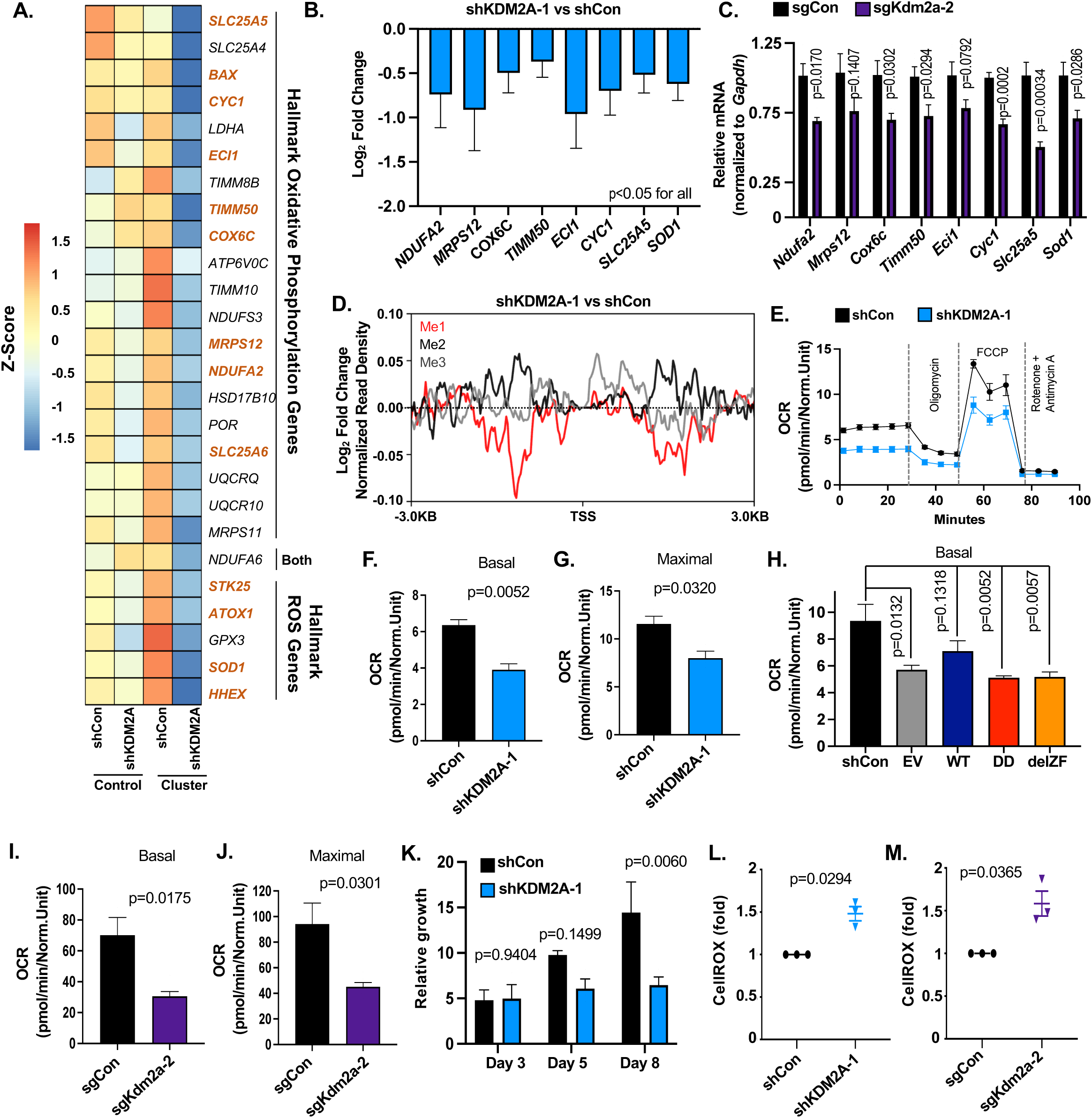
KDM2A demethylase activity sustains an oxidative phosphorylation and redox homeostasis program in metastatic clusters. **(A)** Expression of KDM2A activated genes within the GSEA Hallmark Oxidative Phosphorylation and Hallmark Reactive Oxygen Species pathways was measured by RNA-Seq and plotted for H2030-BrM3 cells under the indicated conditions. Genes were filtered based on unadjusted p-value≤0.05 in shKDM2A-1 vs shCon clusters. Genes in bold are directly bound by KDM2A within 1000bp of the TSS. n=2-3 biological replicates **(B)** Log2 fold change of a subset of KDM2A bound genes from (A). P-value <0.05 for all. N=2-3 biological replicates. **(C)** mRNA expression measured by quantitative polymerase chain reaction (qPCR) in LU-KP cells expressing the indicated sgRNAs for targets from B under cluster forming conditions. P-values by unpaired t-test with Welch’s correction. N=5 biological replicates. **(D)** Log2 ratio of H3K36me1-3 normalized reads in clusters with the indicated shRNAs. Plotted are the average values of N=4 biological replicates. **(E)** Oxygen consumption rate (OCR) was measured after 4 days in H2030-BrM3 clusters with the indicated shRNAs. N=3 biological replicates, each dot represents mean +/− SEM, normalized to cell number. **(F)** Mean basal and **(G)** maximal respiration of samples in E. P-value by Welch’s t-test. **(H)** Mean basal respiration for cell clusters co-expressing shKDM2A-1 with either empty vector (EV), full length KDM2A (WT), 3X KDM2A (DD), or KDM2A (delZF). N=3 biological replicates. P-value by one-way ANOVA with Dunnett’s correction. **(I)** Mean basal and **(J)** maximal respiration of LU-KP clusters with the noted sgRNAs was measured as in (E) after 6 days. P-value by Welch’s t-test. N=6 biological replicates. **(K)** Relative growth of H2030-BrM3 clusters with the indicated shRNAs was measured over the indicated days and quantified as in Figure 1G. P-value by two-way ANOVA with Tukey’s multiple testing. **(L)** Intracellular reactive oxygen species (ROS) was measured using CellROX Deep Red in H2030-BrM3 clusters with the indicated shRNAs after 5 days. Data are normalized to shCon cells and p-value was calculated by one-sided t-test against mean of 1. **(M)** ROS was measured as in (K) for LU-KP clusters expressing the indicated sgRNAs after 5 days. Bar graphs show mean +/− SEM unless otherwise noted.

Major consequences of reduced mitochondrial respiration include diminished cell growth or accumulation of intra-cellular reactive oxygen species (ROS) which can lead to cell death ^44, 45^. Flow cytometric analysis showed that, while cells in clusters proliferate more slowly, no significant difference was observed following KDM2A knockdown (**Extended Data Fig. 5C-F)**. Instead, we observed a decrease in cell viability over time (**Fig. 5K**) which coincided with an increase in mitochondrial ROS (**Extended Data Fig. 5G,H)** as well as total intracellular ROS across independent models (**Fig. 5L,M and Extended Data Fig. 5H**). Additionally, KDM2A inhibition reduced the expression of genes involved in ROS homeostasis (**Fig. 5A-C)**, consistent with a defect in oxidative stress response and cell survival. Therefore, KDM2A H3K36 demethylase activity potentiates the expression of a multi-genic program which promotes the survival of DTC clusters under stress.

### KDM2A inhibition compromises epithelial junctional integrity

Loss of cell-cell junctions promotes epithelial to mesenchymal transition (EMT) and cancer cell dissemination ^46^, and yet the expression of cell junction proteins directly enhances DTC survival and metastatic colonization in some epithelial cancers ^6^. Moreover, perturbation in oxidative phosphorylation can have paradoxical effects on cell migration and cell-cell adhesion ^47^. Therefore, we characterized the effects of KDM2A on the epithelial phenotypes of NSCLC cells. Across NSCLC models that we tested, the clusters appeared epithelial like with the formation of semi permeable tumor cell-cell junctions (**Extended Data Fig. 6A**). H2030-BrM3 clusters express mixed features of epithelial cells (e.g., *CDH1*) and mesenchymal-like cells (e.g., *SNAI1, FN1*) ^46^ (**Extended Data Fig. 6B**). KDM2A knockdown was not sufficient to cause a broad impact on epithelial to mesenchymal plasticity gene expression (**Extended Data Fig. 6B**). However, reduction of KDM2A/Kdm2a across models significantly decreased the expression of epithelial cell junction genes (e.g., *CDH1, JUP, and MARVELD3),* as well as sodium/potassium channel genes (e.g., *KCNK1, KCNK5*) which can regulate cell junction formation ^48, 49^ (**Fig. 6A.B).** Few of these genes are directly bound by KDM2A (**Supplementary Table 4**), suggesting that their repression is a secondary effect of KDM2A inhibition. Knockdown of KDM2A did not affect the migration of H2030-BrM3 cells (**Extended Data Fig. 6C**). Instead, the size distribution of H2030-BrM3 shKDM2A clusters was reduced relative to shCon clusters within 48 hours under cluster forming conditions (**Fig. 6C.D**) and this reduction was most notable after 5-8 days (**Fig. 6E**). Concordant defects in cell aggregation were observed upon Kdm2a knockdown in LU-KP clusters (**Fig. 6F**). These results suggest that KDM2A inhibition diminished the aggregation of metastatic cells. Moreover, this defect phenocopied the effects of treating clusters with the Na+/K+ ATP-ase inhibitor Oubain Octahydrate, which can also prevent epithelial junction formation independently of cell number ^48^ (**Extended Data Fig. 6D,E)**. Immunofluorescence (IF) imaging confirmed that KDM2A/Kdm2a knockdown reduced E-cadherin mediated adherens junctions in clusters as early as 72 hours (**Fig. 6G,H and Extended Data Fig. 6F)**. We conclude that the suppression of KDM2A can compromise epithelial cell junction formation and aggregation of NSCLC clusters.

**Figure 6:**
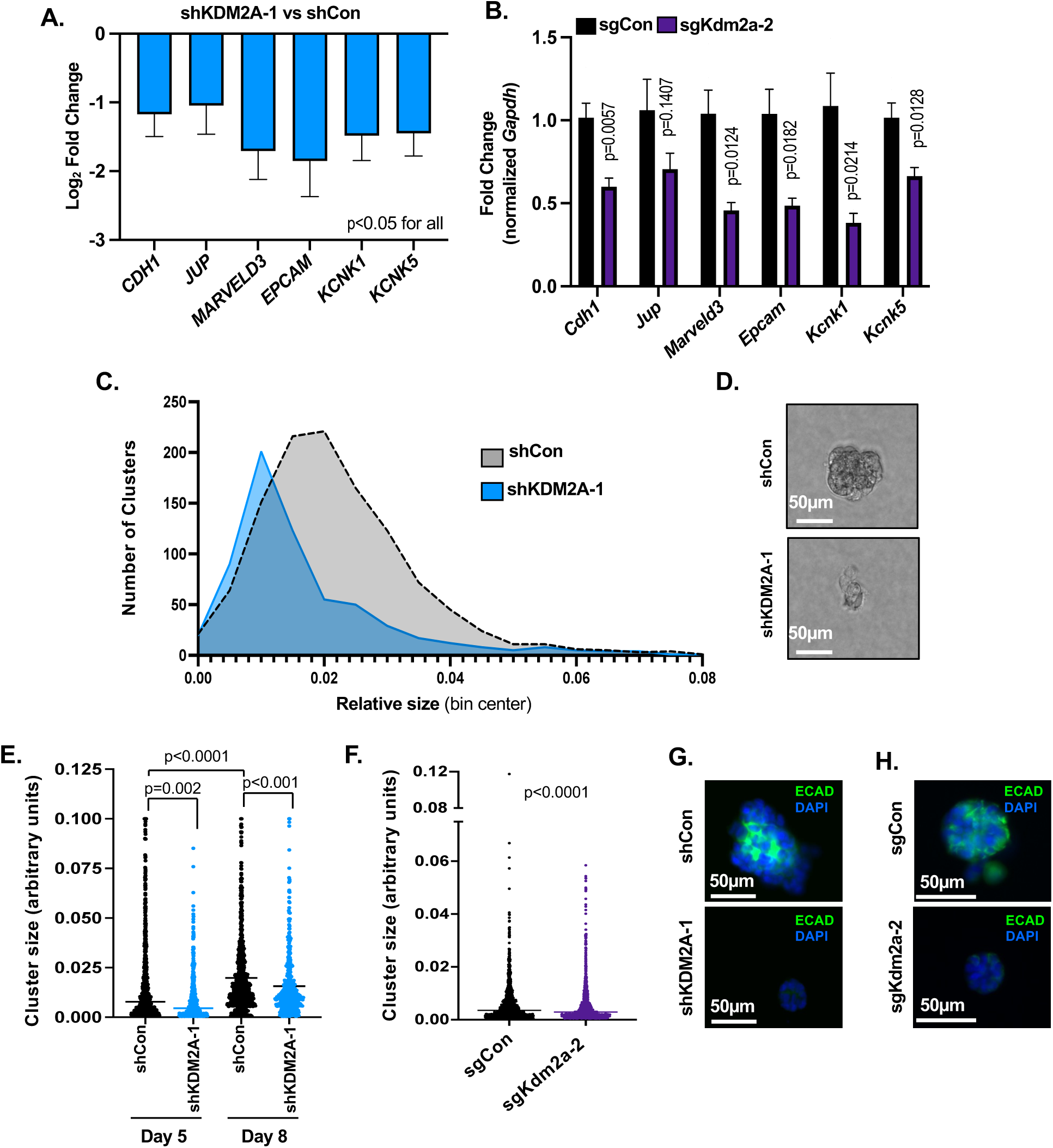
KDM2A inhibition compromises epithelial junctional integrity. **(A)** Log2 fold change of the indicated cell adhesion genes in shKDM2A-1 versus shCon cluster was measured by RNA Seq. p-value <0.05 for all. N=2-3 biological replicates. **(B)** mRNA expression measured by qPCR in LU-KP cells expressing the indicated sgRNAs for genes from (A) under cluster forming conditions. P-values by unpaired t-test with Welch’s correction. n=5 biological replicates. **(C)** Histogram plotting the representative size distribution of H2030-BrM3 clusters, 48 hours after seeding cells. Arbitrary size units were calculated based on RFP positive IF thresholds using Image J. **(D)** Representative bright field images of H2030-BrM3 clusters. Scale=50μm. **(E)** Quantification of the size of H2030-BrM3 clusters expressing the indicated shRNAs at Days 5 and 8. P-values by Kruskal-Wallis test with Dunn’s correction. N=3 biological replicates. **(F)** Quantification of tumor cell cluster size 24 hours after seeding LU-KP cells. Arbitrary size units were calculated based on tdTomato/GFP positive IF thresholds using Image J. P-value by Mann-Whitney. N=3 biological replicates**. (G)** Representative IF images for E-cadherin (green) and DAPI (blue) in H2030-BrM3 clusters with the indicated shRNAs on Day 3. Scale=50μm. **(G)** Representative images of IF staining for E-cadherin (green) and DAPI (blue) in LU-KP clusters with the indicated sgRNAs on Day 3. Scale=50μm.

### KDM2A is required for the seeding and colonization of metastatic cells from circulation

The ability of DTCs to cluster and mitigate metabolic stress in circulation has been associated with poor clinical outcome in human patients with cancer. We therefore examined the expression of KDM2A activated genes in datasets of human circulating tumor cells (CTCs) ^50, 51^. Higher expression of KDM2A activated genes in CTCs from patients with breast cancer correlated with poor survival **(Fig. 7A).** KDM2A activated OXPHOS/ ROS genes (see Fig. 5A) in particular were elevated in those patient derived CTCs found in clusters as opposed to single cell CTCs **(Extended Data Fig. 7A)**. Expression of this signature also correlated with disease progression in CTCs from patients with melanoma (**Extended Data Fig. 7B**). To our knowledge, a similar CTC cohort from human NSCLC is not available for analysis. Alternatively, we confirmed that KDM2A protein is overexpressed in a subset of human primary lung adenocarcinomas (LUAD) ^42^ (**Extended Data Fig. 7C**). *KDM2A* itself is more frequently amplified in human NSCLC brain metastasis as compared to matched primary tumors ^52^. Finally, we found that KDM2A activated OXPHOS/ROS genes were also elevated in metastatic lesions compared to their matched primary tumors from LUAD patients ^18^ (**Fig. 7B and Extended Data Fig. 7D**), suggesting that elevated KDM2A activity could enhance metastatic seeding or colonization in distal sites.

**Figure 7:**
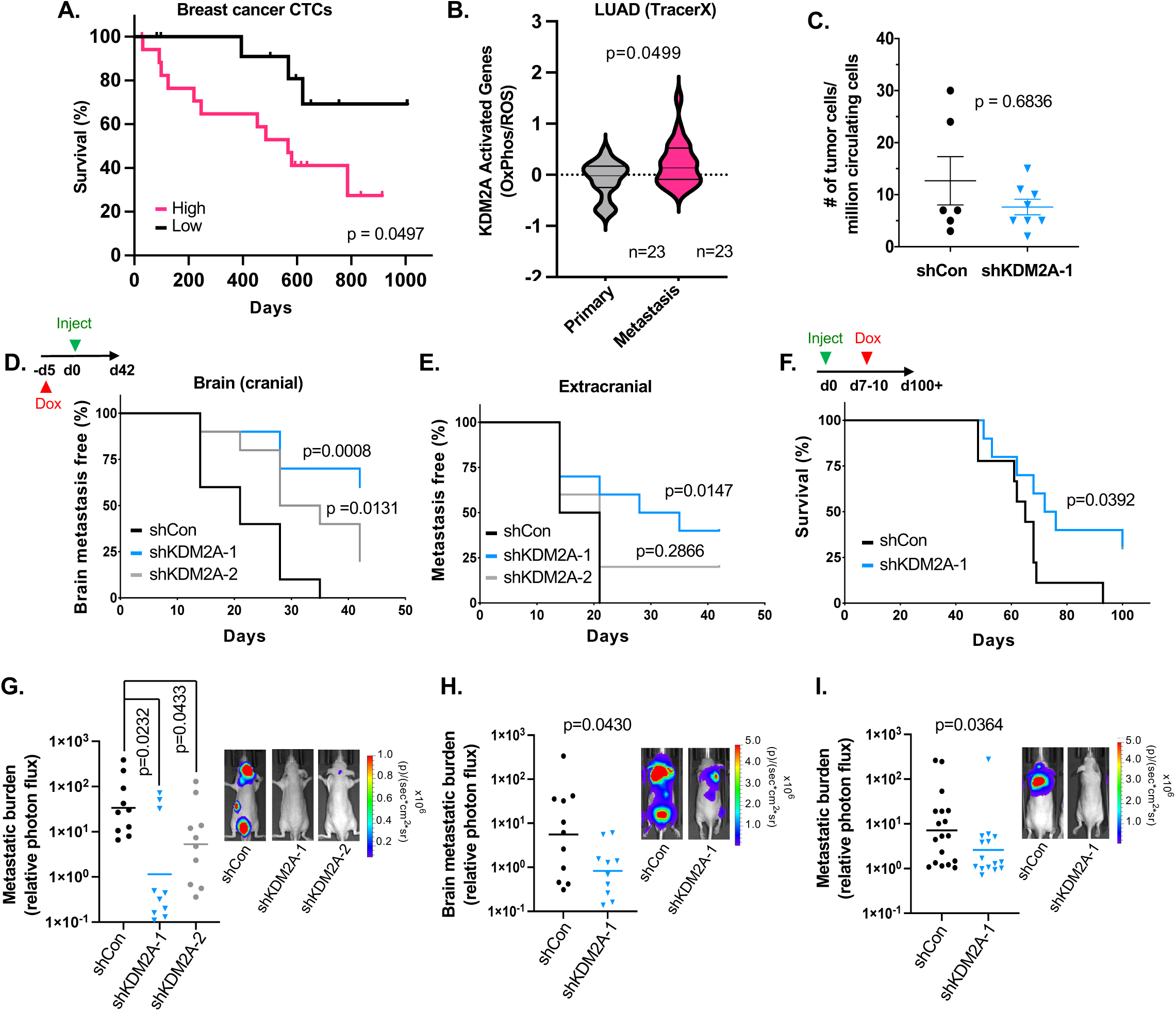
KDM2A mediates metastatic seeding and colonization of tumor cells from circulation. **(A)** Overall survival based on high or low expression of all KDM2A activated genes in the CTCs of breast cancer patients. N=13 low, N=18 high. Raw data are from GSE144495. P-value by Mantel-Cox test. **(B)** Expression of KDM2A bound and activated OXPHOS and ROS genes in RNA-seq samples taken from the TracerX LUAD cohort, comparing primary and metastatic lesions. Raw data are from Zenodo record 10932811. N=23 patients. **(C)** For animals from Figure 3D with median H2030-BrM3 lung tumor burden at Day 42, blood was drawn and RFP positive CTCs detected by flow cytometry. N=6 control; N=8 shKDM2A-1 animals. **(D-E)** H2030-BrM3 cells with the indicated Dox inducible shRNAs were pre-treated with Dox in culture for 5 days before intracardiac injection in athymic mice that were maintained on a Dox containing diet. Metastasis to the brain (D) or extra-cranial sites (E) were detected by BLI and incidence plotted by Kaplan–Meier analysis. P-value by Log-rank test. N=10 animals per group. **(F)** H2030-BrM3 cells with the indicated Dox inducible shRNAs were injected by intracardiac injection into athymic mice and allowed to grow for 7 days before animals were started on a Dox containing diet. Overall survival incidence is plotted. P-value by Log-rank test. N=9 shCon; N=10 shKDM2A-1. **(G)** Endpoint quantification (left) and representative images (right) of metastatic burden as determined by BLI in animals from E-F. **(H)** Endpoint quantification (left) and representative images (right) of brain metastatic burden in animals injected with PC9-BrM4 cells expressing the indicated shRNAs and treated as in (G). N=11 shCon; N=10 shKDM2A-1. **(I)** Endpoint quantification (left) and representative images (right) of metastatic burden in animals injected with MDA-BrM2 cells expressing the indicated shRNAs and treated as in (G). N=19 shCon; N=15 shKDM2A-1. Plots show mean +/− SEM and p-value by Mann-Whitney unless otherwise indicated.

To elucidate which step(s) of metastasis are most susceptible to KDM2A inhibition *in vivo*, we first quantified the number of CTCs in mice after orthotopic injection of H2030-BrM3 cells into the lungs. CTCs were detected in the plasma of mice 42 days after lung tumor growth, consistent with the ability of cancer cells to disseminate from the lungs in this model. Surprisingly however, there was no significant difference in the relative amounts of CTCs between mice with shCon or shKDM2A expressing tumors (**Fig. 7C**), demonstrating that KDM2A is dispensable for intravasation. We and others have shown that H2030-BrM3 cells can extravasate and then seed distant organs as micrometastatic clusters within 7-14 days of entering circulation ^31, 32^. To specifically test the role of KDM2A during metastatic seeding or colonization, shRNAs were induced in H2030-BrM3 cells and then these cells were injected into the arterial circulation of mice. In this experimental setting, knockdown of *KDM2A* significantly decreased the incidence of metastasis in the brain and also extra-cranial sites (**Fig. 7D,E)**. In a separate experiment, we induced KDM2A knockdown *in vivo* 7 days after intra-arterial injection. Inhibition of KDM2A at this delayed time point when cells begin to seed distant sites also impeded metastatic progression (**Extended Data Fig. 7E-G**) and significantly improved overall survival of the animals over 100 days (**Fig. 7F)**.

Finally, given the potential association of KDM2A activity with metastatic outgrowth or poor outcome across cancer types, we compared the effects of KDM2A on metastatic tumor burden in independent models of NSCLC (H2030-BrM3 and PC9-BrM4) and breast cancer (MDA-BrM2), which colonize the brain and other sites after intra-arterial injection ^34, 53^. Inhibition of KDM2A similarly decreased cluster outgrowth in the PC9-BrM4 and MDA-BrM2 models *in vitro* (**Extended Data Fig. 7H,I**). *In vivo,* KDM2A knockdown in the H2030-BrM3 and PC9-BrM4 models abated overall metastatic outgrowth and brain metastasis outgrowth respectively **(Fig.7G,H).** Overall metastatic outgrowth was also reduced in the MDA-BrM2 model **(Fig. 7I)**.

In summary, our *in vivo* data demonstrate that KDM2A activity preferentially increases metastatic seeding and colonization.

## Discussion

Homotypic cell-cell adhesion is a critical mediator of tissue morphogenesis ^27^. Likewise, during metastasis, the formation of cell junctions promotes collective cell migration and enhances the survival of DTCs from epithelial cancers ^54^. The significance of these findings is underscored by studies showing that the detection of CTC clusters in multiple cancers, including NSCLC ^30^, portends poor prognosis. Alterations in histone and DNA modifying enzymes are known to mediate cancer metastasis ^23^. However, the mechanisms by which extracellular cues can specify the epigenome of epithelial cells remain poorly understood. Our study builds on recent findings which intimate that cancer cell aggregation is characterized by dynamic changes in transcription and DNA methylation ^7, 25^. Accordingly, we sought to identify epigenetic dependencies of metastatic cell clusters from NSCLC and revealed that their fitness requires the H3K36 demethylase KDM2A. This contextual requirement for KDM2A was validated across human and murine models with diverse driver mutations.

Dysregulation of H3K36 methylation in human cancer is a significant area of investigation. Oncogenic mutations in *H3K36* prevent H3K36me2/3 methylation in some cancers ^55, 56^. Also, several writers of H3K36 methylation have been implicated in tumor progression ^57^. In NSCLC, increased tumor grade is associated with decreases in H3K36 methylation, due in part to loss of function mutations in the H3K36me3 methyltransferase *SETD2* ^58^. Our observations suggest that, once NSCLC cells disseminate, their metastatic competence is maintained by an H3K36me1 chromatin state that is driven, at least in part, by KDM2A. One limitation of our study is that we did not screen for all potential H3K36 modifiers. It is possible that fluctuations in H3K36me1-3, due to imbalances in H3K36 writer/eraser activity, might also contribute to metastasis through regulation of epithelial lineage plasticity as reported in pancreatic cancer models ^59^. Notably, inhibition of other putative H3K36 erasers, such as KDM2B, may not have a similar impact on epithelial tumor cell clusters, which likely reflects the non-overlapping functions of histone demethylases ^60^. For instance, the N-terminal sequences of KDM2 paralogs are less conserved ^61^. KDM2B preferentially represses transcription and does not require demethylation of H3K36 nor chromatin remodeling to do so ^38, 62^. By contrast, KDM2A, but not KDM2B, forms an acidic patch that binds to nucleosomes and enables access to H3K36me2 residues for demethylation ^63^. Interestingly, *KDM2A* amplification is more frequently detected in NSCLC brain metastasis ^52^, consistent with a distinct role for this H3K36 eraser during metastatic progression.

H3K36me2 is detected across genic and non-coding regions ^64^ and has been correlated with enhancer activation in cancer cells ^65^. It has also been reported that, in NSCLC cells with amplified *KDM2A*, demethylation of H3K36me2 at a few a priori selected genes results in their transcriptional repression ^42^. However, these observations are confounded by the fact that H3K36 methylation is detected across broad stretches of the genome and the molecular consequences of each H3K36 methylation state are likely to be locus dependent. For example, while H3K36me3 marks active genes, this predominantly occurs in gene bodies to repress cryptic transcription and RNA polymerase elongation ^66^. Importantly, H3K36me1, the end product of KDM2 demethylase activity, predominantly occurs in genic regions where it correlates with activated promoters and increased transcription ^64, 67^. There are major gaps in our understanding of KDM2 regulation in cancer because few studies comprehensively assess genome-wide occupancy of KDM2 paralogs in relation to locus specific changes in H3K36 methylation and transcription.

To evaluate the function of KDM2A and H3K36 methylation in metastatic cells, we integrated unbiased epigenome wide analysis with biological experimentation. Our data support a model whereby the primary function of KDM2A is to maintain H3K36 monomethylation at CGI enriched promoters with hypomethylated DNA, thus rendering these loci permissive for transcriptional activation ^68^. This is also consistent with recent studies showing that aberrant H3K36me2 at promoter CGIs leads to *de novo* DNA methylation and transcriptional repression ^69, 70^. It is likely that the sites for H3K36me1/2 modulation are nucleosomes which are accessible to KDM2A when it bound to TSSs. However, in the same biological context, KDM2A could repress transcription at a more limited number of different CGIs, independently of H3K36 methylation. Strikingly, tumor cell-cell adhesion was a determinant of KDM2A recruitment to promoter CGIs, while the expression of KDM2A remained unchanged in tumor cell clusters. Epithelial cell junction formation may induce protein-protein interactions, post-translational modifications of histone demethylases ^71^, and/or altered chromatin architecture that result in KDM2A recruitment to specific promoters. Because DTCs can also aggregate with immune cells ^72^ or endothelial cells ^73^ to form heterotypic clusters, KDM2A activity may also be influenced by stromal cues. Identifying cellular and biochemical determinants of KDM2A transcriptional activity can inform the ongoing therapeutic development of demethylase inhibitors ^74^.

We demonstrated that KDM2A controls the expression of a multi-genic program in tumor cell clusters and that this transcriptional activity correlates with metastasis or poor outcome in patients with various cancer types. KDM2A activated genes included multiple canonical regulators of oxidative phosphorylation. Although tumor initiation and progression are often attributed to increased glycolysis, metastatic cells have an acute dependence on respiratory energy production and expression of oxidative phosphorylation genes as they disseminate ^75, 76^. Consistently, optimal mitochondrial respiration of NSCLC clusters required the CGI binding and JmjC demethylase domains of KDM2A. KDM2A inhibition also caused subsequent disruptions in cell junction integrity, decreased redox homeostasis, and cell survival, which are likely to be collateral effects of dysfunctional mitochondrial respiration. Nevertheless, we cannot exclude the possibility that KDM2A, via transcriptional regulation of different genes, can regulate other cellular functions to enhance metastasis. After cancer cells intravasate, their ability to form clusters increases their intravascular proliferation and survival. Herein, we found that KDM2A is dispensable for the accumulation of cancer cells in circulation but augments their subsequent seeding and colonization of distant organs *in vivo*. Ultimately, inhibiting KDM2A activity decreased metastatic colonization in multiple organs, with a significant impact on brain metastasis, which remains a major clinical challenge in patients with melanoma, breast, and lung cancer ^33^.

In summary, we identified the H3K36 histone demethylase KDM2A as a modulator of tumor cell clustering. We propose that KDM2A binding to chromatin is responsive to cell-cell communication and potentiates the transcriptional activation of multiple genes which are necessary for cellular adaptation during metastasis.

## Methods

### Cell Culture

The H2030-BrM3 ^34^, PC9-BrM4 ^34^, and MDA-MB-231-BrM2 ^53^ are metastatic cell subpopulations of the parental H2030, PC9, and MDA-MB-231 human cell lines, respectively. The aforementioned metastatic subpopulations lines stably co-express GFP and luciferase and were *in vivo* selected for increased metastatic potential in the brain when injected into athymic mice as previously described ^77^. The murine LU-KP cell line was derived from a genetically engineered LSL-Kras^G12D^/p53^fl/fl^ mouse that formed lung tumors after surfactant protein C (SPC) driven Cre recombination ^78, 79^. LU-KP cells were stably transduced to express tdTomato-luciferase (Addgene plasmid #72486). H2030-BrM3, PC9-BrM4, and LU-KP were maintained in treated culture dishes (Corning cat. 430167) and RPMI-1640 (Gibco cat. 11875093) with 10% fetal bovine serum (FBS) (Gibco cat. 10437-028), 1% penicillin/streptomycin (Gibco cat. 15140-122), and 0.2% amphotericin B (Sigma A2942). MDA-BrM2 were maintained as other cell lines but with DMEM (Gibco cat. 11965-092). Cells were regularly tested for mycoplasma contamination using the ATCC Universal Mycoplasma detection Kit (ATCC cat. 30-1012K).

For adherent sub-confluent control conditions, ∼5×10^2^ tumor cells per cm^2^ of surface area were seeded in treated culture dishes as above, but with 2% FBS. For cluster forming conditions, cells were seeded in 2% FBS in ultra-low attachment plates (Corning cat. 4615, 3261, 3474, 3471, 3743) and supplemented with 5% growth factor reduced (GFR) Matrigel extracellular matrix (Corning cat. 356231). Media was refreshed every 3-4 days. Where indicated, cultures were treated with KDM2A/7A-In-1 (CAS No. 2169272-46-0, MCE cat. HY-108706) and ouabain octahydrate (CAS No. 11018-89-6, MCE cat. HY-B0542) at the indicated concentrations. Doxycycline and drugs where used were replenished with media every 3-4 days.

### shRNA, sgRNA, and cDNAs

Doxycycline (Dox) inducible shRNA cell lines were generated in the pINDUCER10 ^80^ backbone. Dox inducible sgRNA cell lines were generated in in the TLCV2 ^81^ backbone. Viruses were generated as previously described using 4^th^ generation packaging constructs, ^35^ titered, and cells infected at a multiplicity of infection (MOI) of 1. Cells were selected using 2µg/mL of puromycin (Gibco cat. A1113803). For shRNA induction, cells were treated with PBS or Dox at 0.5 µg/mL. For sgRNA expression, LU-KP were treated with 0.5 µg/mL for 7 days to induce Cas9 activity. Cas9-GFP positive cells were isolated by fluorescence activated cell sorting to generate LU-KP cell lines with stable knockdown. shRNA and sgRNA constructs used are shown below.

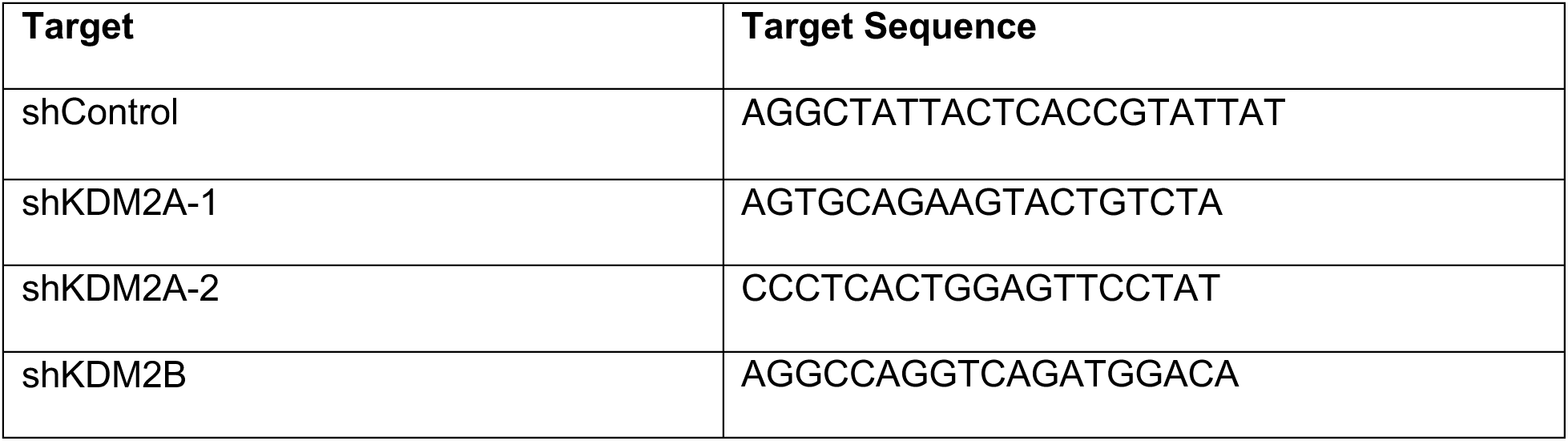

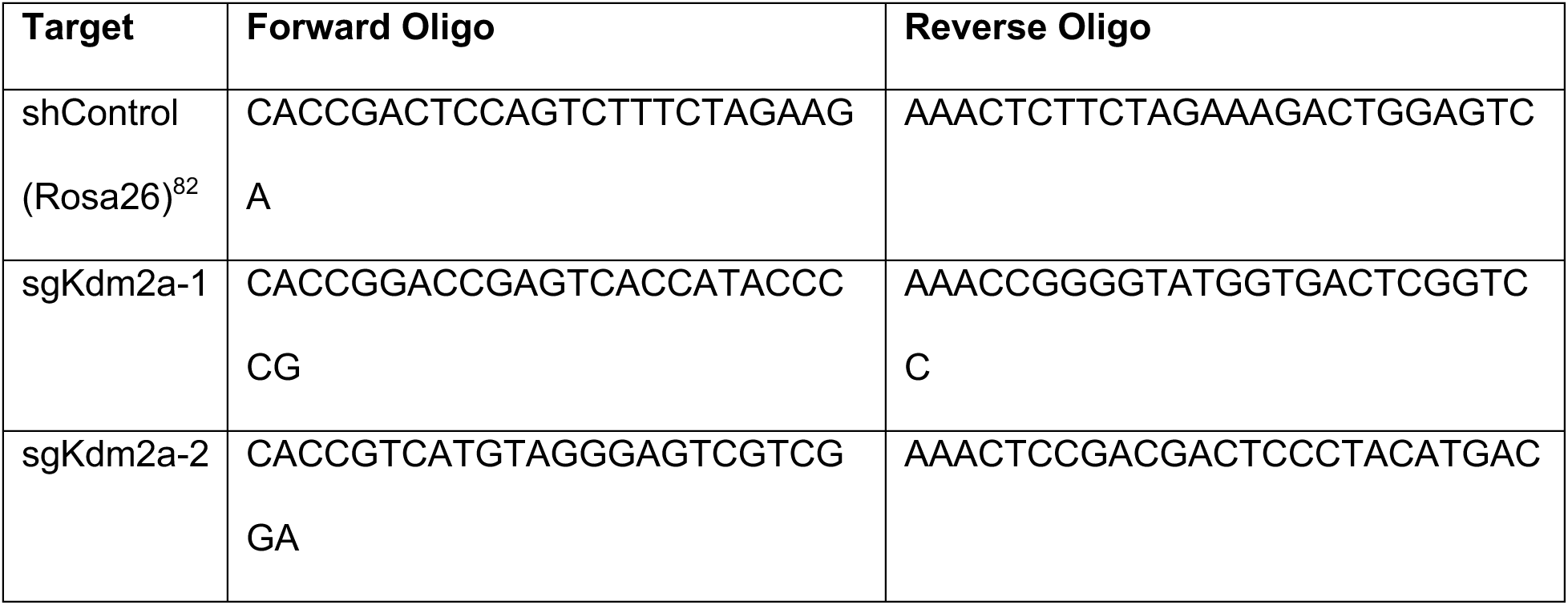

To ectopically rescue expression of full length KDM2A or its mutants, a FLAG tag (DYKDDDDK) sequence was subcloned in triplicate next to the N-terminus of the full length *KDM2A* coding sequence or the demethylase deficient ^36, 37, 83, 84^ (DD) or zinc finger deletion ^37, 84, 85^ (delZF) *KDM2A* mutants. cDNAs were subcloned into the PInducer20 lentiviral backbone with blasticidin as a selectable marker ^80^. Silent mutations of the targeted sequence: (AGTGCAGAAGTACTGTCTA) to (AGTTCAAAAATACTGCCTG) were introduced to confer resistance to shKDM2A-1 mediated knockdown. PInducer20 with empty vector or expressing cDNAs were then used to infect H2030-BrM3 cells expressing shCon or shKDM2A-1. All rescue cell lines were selected with blasticidin (10ug/ml) and maintained in Dox (0.5 µg/mL) throughout. **shRNA screen**. An inducible, lentiviral barcoded short hairpin RNA (shRNA) library targeting 89 epigenetic regulators was previously assembled and validated for knockdown efficiency and *in vitro* screens in metastatic cells ^35, 86^. In brief, lentiviruses encoding each shRNA were used to infect H2030-BrM3 cells (MOI=1), which were then selected with puromycin. 40-45 individually infected cell lines (including 3 lines expressing positive or negative control shRNAs) were combined at equal cell numbers into minipools. Minipools were cultured under control or cluster forming conditions, +/− dox, for 12 days. Samples were harvested for genomic DNA extraction and qPCR was performed to determine the abundance of each barcode relative to day 0. The TRE element present in each shRNA expressing construct was used as a housekeeping control. **Animal Studies**. All animal work was conducted in accordance with the Yale University Animal Care and Use Committee guidelines. For subcutaneous studies, 5×10^5^ cells were suspended in a 1:1 mixture of sterile PBS and complete Matrigel (Corning cat. 356237) and injected into a single flank of male C56BL/6J mice from The Jackson Laboratory (strain code 664, aged 5-6 weeks). Tumors were measured by digital caliper at least 2x/week, and tumor volume was calculated using the formula (vol=π*length*width^2^/6). At the first time point at which tumor volume exceeded 0.7cm^3^, animals were euthanized. Lungs were collected and imaged using an IVIS Spectrum (PerkinElmer). Lung tumor burden was quantified as the number of independently detectable sites above a shared minimum baseline flux.

For intracardiac injections, cells were dissociated and resuspended into sterile PBS and kept on ice. 5×10^4^ cells/100µL were injected into the left ventricle of the heart of male NCI athymic nude mice (NCI strain code 553, aged 5-6 weeks) purchased from Charles River Laboratories. For injection of clusters into mice, cells were cultured overnight at 5×10^4^ /100µL in cluster forming conditions (24-wells). To ensure consistency one well was prepared for each animal. Prior to injection, each well was collected and gently spun down (5 minutes, 1500rpm). Clusters were washed and resuspended in PBS. For intratracheal injections, lungs were pre-conditioned with bleomycin for two weeks before injecting 2.5×10^4^ tumor cells into the trachea as previously described ^87^. Animals were maintained on Dox chow throughout the course of the experiment. Tumor outgrowth was measured weekly through bioluminescent imaging using either an IVIS Spectrum or IVIS X5 (PerkinElmer). Animals were sedated with isoflurane and given 0.1 mL luciferin (Perkin Elmer cat. 122799) retro-orbitally prior to imaging. For imaging of tissue at necropsy, animals were given 0.2 mL luciferin directly prior to euthanasia. For shRNA knockdown *in vivo*, animals were switched to a doxycycline chow diet at the indicated time points (Envigo, cat. TD.01306).

### Clonogenic growth

1×10^3^ cells were seeded in triplicate in 6-well tissue culture plates. Media was changed from 10% FBS to 2% FBS 24 hours after seeding. After 7-14 days, plates were washed 1x with PBS and fixed for 10 minutes at room temperature (RT) in 4% paraformaldehyde in PBS. Fixed colonies were washed 2x with PBS and incubated in 0.05% crystal violet for 1h at RT. Plates were then rinsed in ddH_2_O and imaged by Bio-Rad ChemiDoc MP Imaging system. Colony area was quantified in Fiji (ImageJ) using the ColonyArea package ^88, 89^.

### Cluster Outgrowth

1×10^3^ cells were seeded in 24-well ultra-low attachment plates under cluster forming conditions. For all experiments, quadruplicate samples were collected at day 0. At endpoint, samples were lysed with Passive Lysis Buffer (Promega cat. E1941), and cell number was estimated based on the luciferase assay system (Promega cat. E1500). Growth was calculated by normalizing luciferase levels at the indicated time points to day 0 levels.

### Western Blots

Cells were washed 1x with ice cold PBS before lysis in RIPA buffer. Clusters were spun down (1500rpm, 5 minutes, 4°C) prior to washing and lysis. Samples were incubated on ice for 15 minutes with inversion every 5 minutes, then spun for 15 minutes (13,000 RPM, 4°C). Supernatant was collected and quantified via Bradford assay (Bio-Rad cat.5000201). 10-50 µg of protein was loaded into gels prior to electrophoresis and transferred onto nitrocellulose membranes (Bio-Rad cat. 1620112) using the Mini-PROTEAN Tetra Cell (Bio-Rad cat. 1658001EDU). Membranes were blocked in TBST containing 5% non-fat milk for 1 hour and then incubated with primary antibody overnight in the same. Primary antibodies are listed in the table below. Membranes were washed 3x for 10-15 min with TBST and incubated for 1h with secondary antibody at 1:2500 dilution for rabbit (Invitrogen cat. 31460) or 1:3000 dilution for mouse (Invitrogen cat. 31437). Membranes were washed 3x for 10-15 minutes before development with SuperSignal Pico/Femto PLUS Chemiluminescent Substrate (Thermo Scientfic cat. 34577 and 34094). All blots were imaged using the Bio-Rad ChemiDoc MP Imaging system and processed in Image Lab Software (Bio-Rad).

| Antibody | Company | Catalog # | Target Species | Dilution |
| --- | --- | --- | --- | --- |
| KDM2A/Kdm2a | Abcam | ab191387 | Rabbit | 1:1000 |
| FLAG | Sigma-Aldrich | F1804 | Mouse | 1:1000 |
| E-Cadherin | Cell Signaling | 4650S | Rabbit | 1:1000 |
| Tubulin | Sigma-Aldrich | T5168 | Mouse | 1:1000 |

### Immunofluorescence

Cells from adherent cultures were washed with PBS before being fixed with 4% paraformaldehyde. Clusters were collected by mixing media 1:1 with ice cold PBS and spinning onto chambered slides (Thermo Scientific cat. 154534PK) (1500rpm, 5 minutes, 4°C). Slides were washed with ice cold PBS and spun before fixation with 4% paraformaldehyde. Cells were permeabilized with 0.5% Triton X-100 in 3% bovine serum albumin (Sigma Aldrich cat. A9647) for 1h at RT for KDM2A and Streptavidin staining, and 0.1% Triton X-100 in PBS for 10 minutes for other proteins listed below. Samples were blocked for 1h at RT with 10% goat serum. Secondary staining and amplification were performed using Alexa Flour Tyramide SuperBoost kits (Invitrogen cat. B40926, B40912) according to manufacturer recommendations. All samples were mounted in Prolong Gold Antifade Reagent with DAPI (Invitrogen cat. P36935). For junctional staining, cells were labeled for 30 minutes at 37°C with 0.8mM NHS-Biotin (Thermo Scientific cat. A39256) in PBS prior to collection and incubating with 1:200 Streptavidin-FITC (Invitrogen SA1000) for 2-3h at room temperature.

| Antibody | Cell Line | Company | Catalog # | Target<br>Species | Dilution |
| --- | --- | --- | --- | --- | --- |
| KDM2A | H2030-BrM3 | Abcam | ab191387 | Rabbit | 1:200 |
| E-Cadherin | H2030-BrM3 | Abcam | ab231303 | Mouse | 1:200 |
| ZO-1 | H2030-BrM3 | Abcam | ab221597 | Rabbit | 1:200 |
| E-Cadherin | LU-KP | Cell<br>Signaling<br>Technology | 3195S | Rabbit | 1:200 |

### Cluster size

Clusters were imaged on a Keyence microscope using phase contrast microscopy and fluorescent detection where possible. Analysis of fluorescent images was semi-automated using a Fiji (ImageJ) macro to open, 8-bit convert, threshold, and measure cluster size using the “Analyze Particles” tool (ImageJ)^88^. Whole wells or representative regions were sampled for imaging. Non-fluorescent measurement was done on phase contrast images in Fiji using the freehand selection tool. Arbitrary units reflect image sizes used for quantification and are consistent within each experiment. Measurements were compiled and plotted in GraphPad Prism.

### Quantitative Real Time PCR (qPCR)

Samples were washed with PBS prior to lysis in RLT buffer, homogenized using QIAShredder columns (Qiagen cat. 79656), and RNA extracted with the RNeasy kit (Qiagen cat. 74016). Synthesis of cDNA was done using the iScript kit (Bio-Rad cat. 1708891) and diluted 10-fold prior to use. RT-qPCR reactions were run using SYBR chemistry using iTaq Universal SYBR Green Supermix (Bio-Rad cat. 1725124) on a Viia7 (Applied Biosystems) or QS5 (QuantStudio). All reactions were assayed in quadruplicate with normalization to housekeeping genes *GAPDH/Gapdh* or *HPRT1*, and quantification determined by ΔΔCt approach. A table of primer sequences is below.

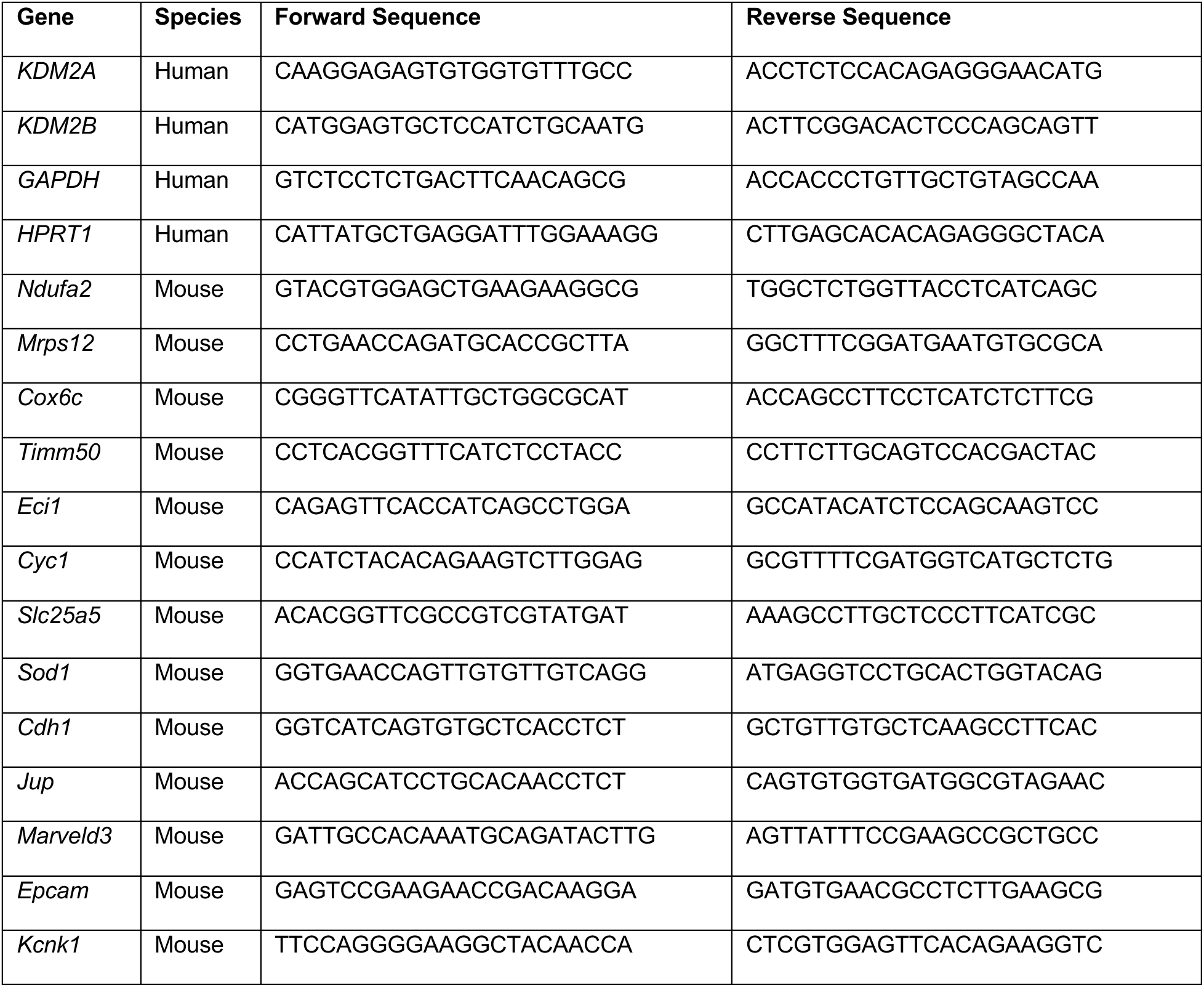
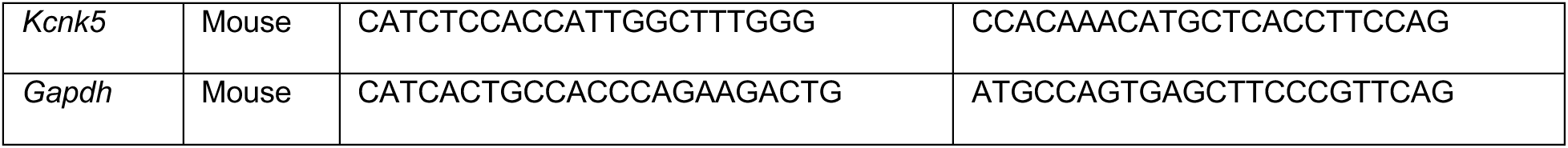

### RNA-sequencing

RNA was collected as described above with the addition of a 15 minute on-column DNAse treatment (DNAse Qiagen cat. 79254). RNA quality was confirmed by Agilent Bioanalyzer, and TruSeq PolyA enriched libraries were generated by the Yale Stem Cell Center Genomics Core. Samples were sequenced on an Illumina HiSeq 4000 to generate a minimum of 25M paired-end reads. Read quality was checked using FastQC (Babraham Bioinformatics), adaptors trimmed with TrimGalore (Babraham Bioinformatics), and reads were aligned to hg38 using HISAT2^90^. Reads were filtered using Picard Tools (Broad Institute), transcripts assembled using String Tie with Gencode v27, and counts generated using Ballgown ^91^. Differential expression was determined using DESeq2 ^92^. Pathway analysis was conducted using Gene Set Enrichment Analysis (GSEA) on all detected transcripts (no significance threshold filtering) ^93^. Z-scores were generated for genes of interest from normalized count data in R using the formula ((sample expression of gene - mean expression of gene across all samples)/ standard deviation of gene expression across all samples). Heatmaps of z-scores were plotted using the R package pheatmap. Genes inferred to be activated or suppressed by KDM2A were generated based on log_2_ fold changes between shKDM2A-1 and shControl clusters.

### Chromatin Immunoprecipitation and Sequencing (ChIP-Seq)

FLAG-KDM2A samples were double-fixed using 2mM ethylene glycol bis(scuccinimidyl succinate) (EGS) (Thermo Scientific cat. 21565) for 1h followed by 1% formaldehyde for ten minutes. H3K36 methylation samples were fixed with 1% formaldehyde alone. Reactions were quenched with glycine to a final concentration of 0.125M. Samples were washed with PBS and lysed in complete sonication buffer (20mM Tris pH 8.0, 2mM EDTA, 0.5mM EGTA, 1x protease inhibitors (Roche), 0.5% SDS, and 0.5mM PMSF) prior to sonication with a QSonica Q800. Fragment size (200-500bp) from extracted DNA was confirmed by agarose gel and quantified by Qubit. Input was aliquoted from 20% of total material, and chromatin samples were subsequently pre-cleared for 1 hour at 4°C with immunoprecipitation (IP) buffer (20mM Tris pH 8.0, 2mM EDTA, 0.5% Triton X-100, 150mM NaCl, and 10% Glycerol) and Protein A/G beads (Thermo Scientific cat. 20421). IP for proteins of interest was conducted using protein A/G beads and 10µg of antibody (see table below). For H3K36 methylation IPs Drosophila spike-in chromatin (100ng) (Active Motif cat. 53083) and spike-in antibody (5μg) (Active Motif cat. 61686) were added. IPs were washed then eluted in a buffer of 100mM NaHC03 and 1% SDS. Both IP and input samples were proteinase K treated, reverse crosslinked, and purified by phenol-chloroform extraction with ethanol precipitation. Quality control and library preparation was done by Yale Stem Cell Center Genomics Core. In brief, samples were checked by Agilent Bioanalyzer, and libraries were prepared by the using ThruPlex DNA Seq (Takara Bio. cat. R400674) for H3K36 libraries, and DNA SMART ChiP-Seq (Takara Bio.cat. 634865) for FLAG libraries. Libraries were barcoded, pooled, and sequenced (PE) on Illumina NovaSeq X and NovaSeq 6000 machines.

H3K27Ac immunoprecipitation, library preparation, and sequencing was conducted commercially by Active Motif (Carlsbad, CA). In brief, cells were fixed with 1% formaldehyde for 15 min and quenched with 0.125 M glycine. Chromatin was isolated by adding lysis buffer, followed by disruption with a Dounce homogenizer. Lysates were sonicated and the DNA sheared to an average length of 300-500 bp with an EpiShear probe sonicator (Active Motif cat. 53051). Genomic DNA (Input) was prepared by treating aliquots of chromatin with RNase, proteinase K and heat for de-crosslinking, followed by SPRI beads clean up (Beckman Coulter) and quantitation by Qubit. Extrapolation to the original chromatin volume allowed determination of the total chromatin yield. 30 µg of chromatin was precleared with protein A agarose beads (Invitrogen), and regions of interest were isolated using 4ug of antibody against H3K27Ac. Complexes were washed, eluted from the beads with SDS buffer, and subjected to RNase and proteinase K treatment. Crosslinks were reversed by incubation overnight at 65 C, and ChIP DNA was purified by phenol-chloroform extraction and ethanol precipitation.

Custom type Illumina sequencing libraries were prepared from the ChIP and Input DNAs on an automated system (Apollo 342, Wafergen Biosystems/Takara). After a final PCR amplification step, the resulting DNA libraries were quantified and sequenced on a NextSeq 500 (Illumina).

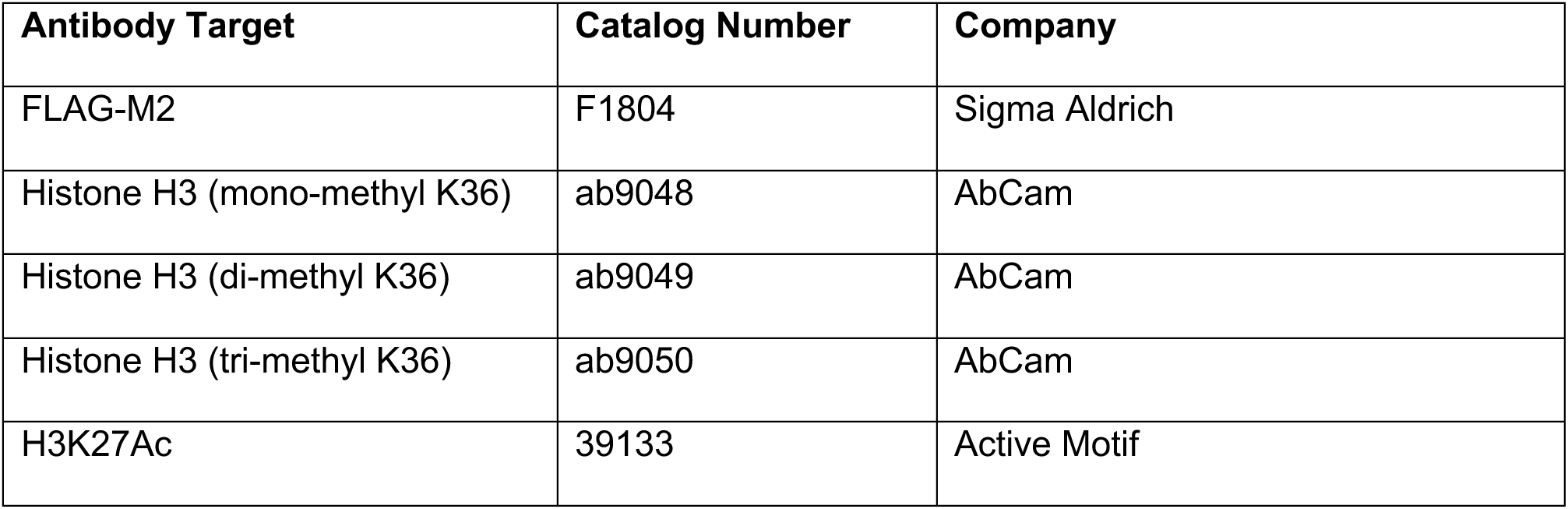

### ChIP-Seq Analysis

All samples were quality controlled using FastQC (Babraham Bioinformatics). Narrow peaks (H3K27Ac, FLAG-KDM2A) were trimmed with TrimGalore (Babraham Bioinformatics) and FLAG-KDM2A reads were cleaned with SMARTCleaner ^94^. Broad peaks (H3K36me) were trimmed Trimmomatic ^95^. All experiments were aligned against hg38 using Bowtie2 ^96^. Narrow peak calling was done using MACS2 and broad peak calling with MACS3^97^ using the -broad flag. Blacklisted regions, PCR optical duplicates, and poor-quality reads were filtered in all cases. Peaks were annotated and motifs identified using HOMER ^98^ “annotatePeaks” and “findMotifs” functions using both hg38 and Gencode v27 as inputs.

Coverage files were generated using deepTools ^99^ tools bamCoverage, bigwigAverage, and bigwigCompare. Prior to coverage file generation, optical duplicates were removed using Picard Tools (Broad Institute), and poor-quality reads and blacklisted regions filtered using samTools ^100^. All coverage files were generated using bamCoverage with normalization by the counts per million (CPM) method, and average coverage of replicates was generated using bigwigAverage. Log_2_ ratio tracks of experimental versus control conditions were generated using bigwigCompare. Coverage tracks were plotted over custom bed files of interest generated using the UCSC Table Browser ^101^. Data was visualized and quantified using the computeMatrix, plotProfile, and plotHeatmap functions of deepTools. Quantification was exported from underlying matrixes and plotted in GraphPad Prism.

### Analysis of sc-RNA and bulk RNA sequencing in patient biospecimens

Single cell RNA-seq data was obtained from Hong et al.^51^ (GSE157743) and Ebright et al.^50^ (GSE144495) and bulk RNA-seq from Martinez-Ruiz et al. ^18^ (TracerX consortium, Zenodo 10932811). Single cell count data was normalized using Linnorm ^102^ with default settings and DataImputation=TRUE, while VST transformed bulk RNA-seq data was extracted using a modification of the script ““6_top_fig4_edfig4.R” from the original manuscript. Samples were further z-score normalized on a per-gene level, before calculation of the mean expression of KDM2A activated genes for each cell or tissue sample. Where applicable, we classified patients into high or low signature expression based on median differential expression of the signature across the cohort. For survival curves, patients with multiple CTCs were stratified based on the average signature expression of their samples. Likewise, for patients with more than one primary or metastatic site sampled in the bulk RNA-seq, average expression of those sites was used. Primary tumor versus normal tissue protein comparisons were plotted using Normalized Relative Protein Expression from Protein Atlas (PDC000153). Processed data was plotted and statically analyzed using GraphPad Prism.

### Flow Cytometry Analysis

To assay proliferation, cells were labeled for 4 hours with 10 µM BrdU (Invitrogen cat. B23151) at 37°C in culture medium. Adherent cells were harvested and clustered cells dissociated by treatment with 0.25% trypsin, and fixed in 4% paraformaldehyde in PBS for 20 minutes at RT. After fixing, cells were permeabilized with 0.5%Triton-X 100 in PBS for 15 minutes at RT, DNAse treated, and incubated for 20 minutes with anti-BrdU antibody (Abcam cat. ab22075) in staining buffer (PBS w/o Ca^2+^ or Mg^2+^, 1% FBS, 0.09% NaN_3_). Cells were washed 2x between each step and stained with DAPI for 5 minutes directly prior to analysis on the BD LSR-II. For ROS staining, cells were incubated with CellROX Deep Red dye (Invitrogen cat. C10491) or MitoSOX Green dye (Invitrogen cat. M36006) for 30 minutes at 37°C, washed, and run on a BD LSR-II. All results were analyzed in FloJo.

### Mitochondrial Stress Test

Yale Chemical Metabolism Core employed a Seahorse XF Analyzer (Agilent). Cells were seeded at a density of 2.5×10^4^ cells/well in 96-well Seahorse XF Plates (Agilent cat. 103794-100) 24 hours prior to the assay. Cells were washed 1x with then switched to assay medium (DMEM, 11mM glucose, 2mM glutamine, 5mM HEPES, 0.2%) prior to the experiment. Clusters were spun down (5 min, 1500 rpm), washed 1x in assay medium, and spun onto 96-well Seahorse XF Plates directly before analysis. Five baseline measurements were taken/experiment, followed by sequential injection of Oligomycin (1 µM), FCCP (3 µM for H2030 BrM3 cells/5 µM for LU-KP cells), and Rotenone/Antimycin A (5 µM/10 µM) with three additional measurements after each treatment. At least 6 technical replicates were included per condition. Cell number was estimated by Hoechst or luciferase, and results were normalized per cell number.

### Quantification and Statistics

Data were plotted and analyzed in GraphPad Prism (version 10.5.0) unless otherwise noted. Graphs show mean ± SEM unless otherwise noted. Number of biological replicates is denoted in figure legends using “n=”, with a minimum representation of 2. Parametric tests were not used unless samples passed normality testing, and multiple testing corrections applied where appropriate. Standard deviations and population distributions were examined prior to test selection.

### Data Availability

RNA-seq and ChIP-seq data generated for this manuscript were deposited at GEO under accession number GSE316477 and GSE316752. Clinical datasets were obtained from the publicly available accession numbers GSE157743, GSE144495, Zenodo 10932811, and PDC000153.

## Author Contributions

Conceptualization: C.K., K.D.P., E.W., and D.X.N; Methodology: C.K., K.D.P., T.F.W.; Investigation: C.K., K.D.P., E.W., M.Z., S.C., T.T., Y.H.; Analysis: C.K., K.D.P., D.Z., Q.Y., and D.X.N.; Writing: C.K., K.D.P., Q.Y., and D.X.N.; Supervision: D.X.N.

## Acknowledgements

This work was funded by the National Cancer Institute (NCI) and Department of Defense including: R01CA166376 (D.X.N.), P50CA196530 (D.X.N).; HT9425-23-1-0602 (Q.Y.), HT9425-23-1-0603 (D.X.N.), T32CA193200 (C.K), F31CA261126 (C.K.), as well as the Beatrice Kleinberg Neuwirth Fund (D.X.N.). Additional technical support was provided by the Yale Center for Genome Analysis, Yale Cancer Center Shared Resources (P30CA016359), Yale Flow Cytometry, Yale Stem Cell Center, and Yale Chemical Metabolism Core.

---

**Extended Data Figure 1:**
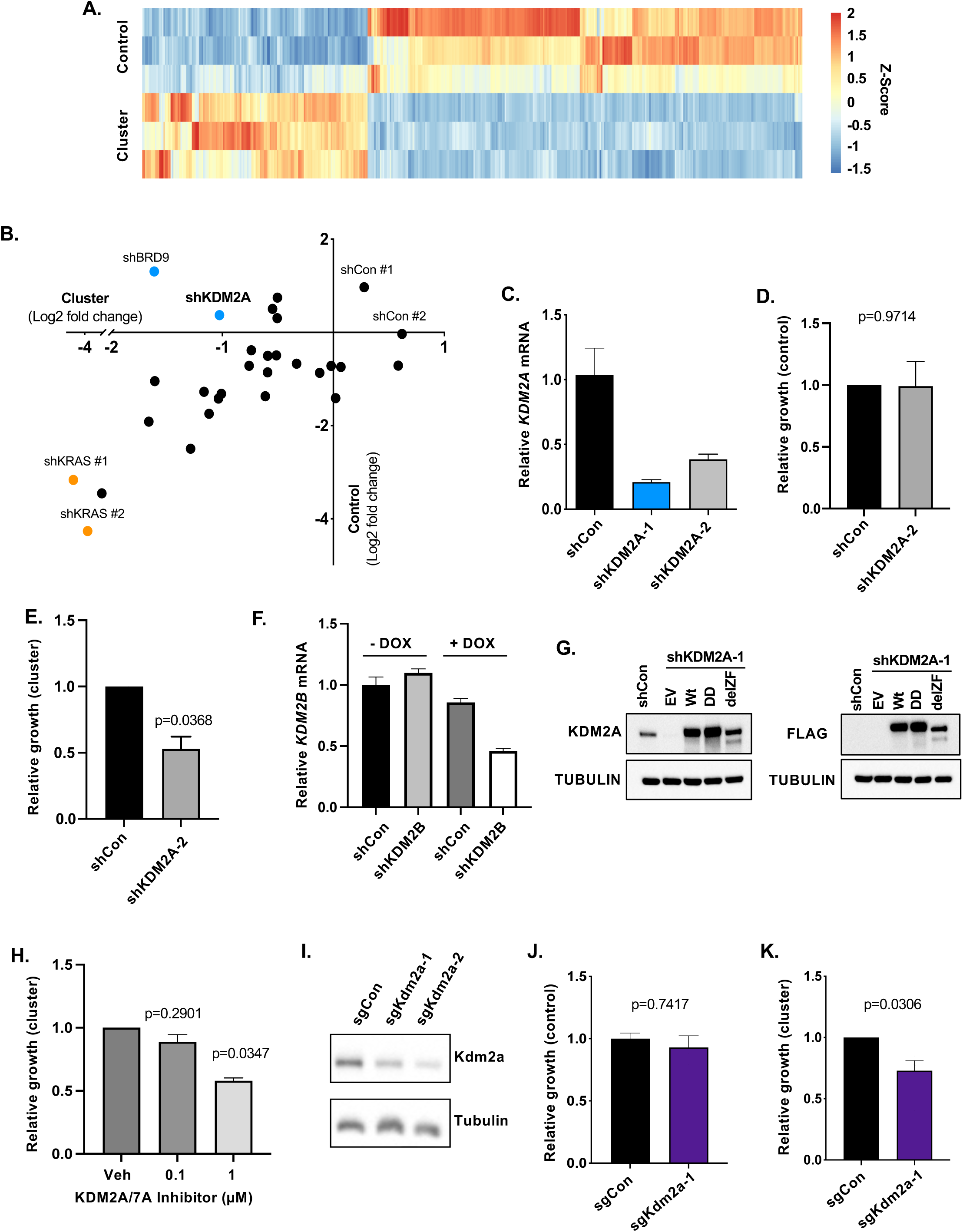
Metastatic cell clusters are dependent on KDM2A. **(A)** Bulk RNA-sequencing was performed on H2030-BrM3 cells under sub-confluent (control) or cluster forming conditions (cluster) after 5 days. Heatmap depicts Z-score transformed levels of the top differentially expressed genes (n=466). **(B)** Plot summarizing fold change of barcode expression in pooled H2030-BrM3 cells under cluster forming (x axis) versus control (Y axis) conditions. **(C)** *KDM2A* mRNA was measured by quantitative polymerase chain reaction (qPCR) in H2030-BrM3 cells expressing the indicated shRNAs and treated with Dox for 3 days. Data are normalized to *HPRT1* expression. N=4 technical replicates. Bar graphs show mean fold change relative to shCon with standard deviation (SD)**. (D)** Clonogenic growth of H2030-BrM3 cells expressing shCon or an independent shRNA targeting *KDM2A* (shKDM2A-2) measured after 10 days as in Figure 1F. P-value by one-sample t-test against mean of shCon. N=2 biological replicates. **(E)** Relative growth of H2030-BrM3 tumor cells expressing the indicated shRNAs under cluster forming conditions as in Figure 1G. P-value by one-sample t-test against mean of shCon samples. N=3 biological replicates. **(F)** *KDM2B* mRNA was measured as in (C) for cells expressing the indicated shRNAs with or without Dox for 5 days. Results are normalized to *GAPDH* expression. N=4 technical replicates. Bar graphs show mean fold change relative to shCon with SD**. (G)** Western blot of endogenous KDM2A (left) and FLAG tag (right) in H2030-BrM3 cells co-expressing shKDM2A-1 and one of the indicated 3X FLAG-KDM2A cDNAs. Cells were treated with Dox for 5 days. Tubulin was used a loading control. **(H)** Relative growth of H2030-BrM3 cell clusters treated with a KDM2A/7A inhibitor at the indicated doses measured after 12 days as in (E). P-value by one-sample t-test against mean of vehicle control. N=2 biological replicates. **(I)** Western blot for Kdm2a in LU-KP cells expressing the indicated sgRNAs after Dox treatment for 7 days. Tubulin was used a loading control. **(J)** Clonogenic growth of LU-KP cells expressing sgCon or sgKdm2a-1 was quantified as in Figure 1J. P-value by following one-way ANOVA with Dunnett’s multiple testing correction against both sgKdm2a-1 and sgKdm2a-2 from Figure 1J. N=3 biological replicates. **(K)** Outgrowth for LU-KP cluster with the indicated sgRNAs was measured as in Figure 1K. N=5 biological replicates. P-value by one-sided t-test against mean of sgCon (same samples as in Figure 1K). Bar graphs show mean +/− SEM unless otherwise noted.

**Extended Data Figure 2:**
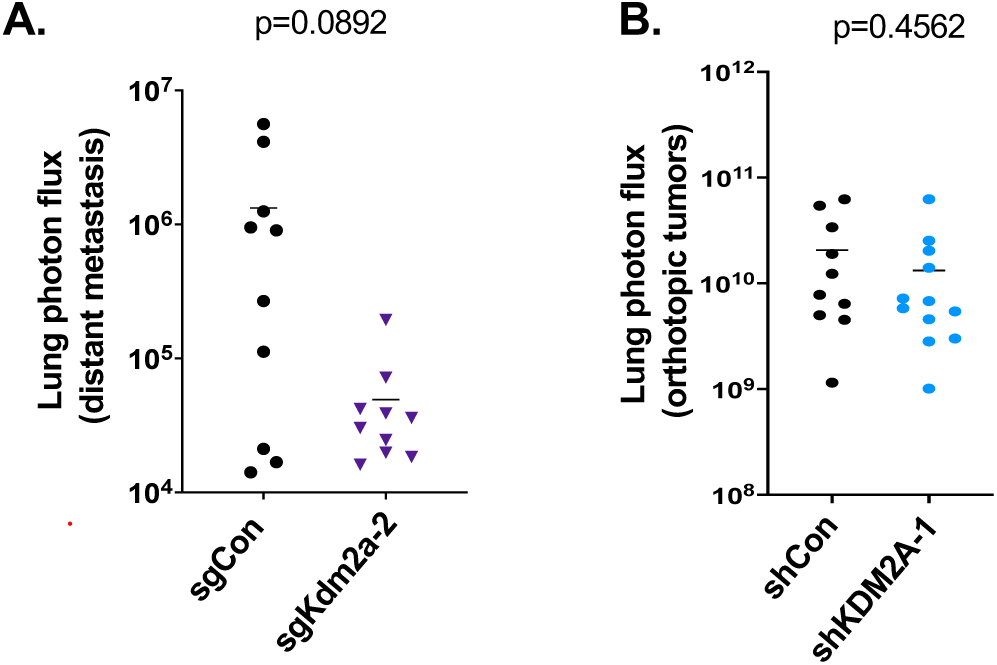
KDM2A mediates spontaneous NSCLC metastasis. **(A)** Metastatic burden was measured by BLI for lungs with metastases quantified in 2C. P-value by Mann-Whitney. **(B)** Orthotopic lung tumor burden was measured by BLI for animals from Figure 2E. P-value by Mann-Whitney.

**Extended Data Figure 3:**
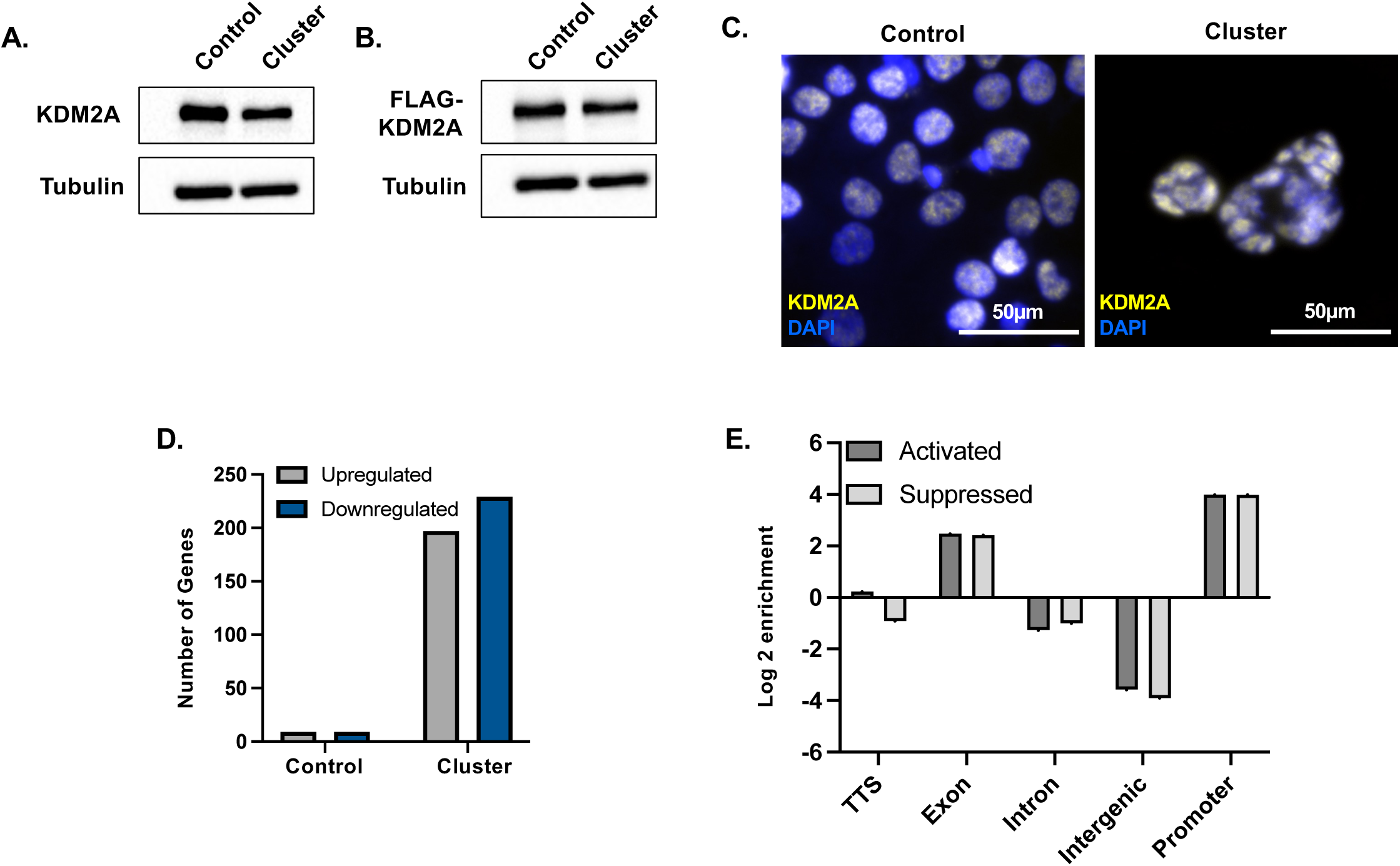
Effects of tumor cell aggregation on KDM2A localization, chromatin binding, and transcription. Western blot of endogenous KDM2A **(A)** and **(B)** FLAG-KDM2A levels between control and clustered H2030-BrM3 cells. **(C)** IF of endogenous KDM2A localization in H2030-BrM3 cells cultured as in (A). **(D)** Number of all genes (KDM2A bound and unbound) that are differentially expressed between shCon and shKDM2A-1 H2030-BrM3 cells, grown under control or cluster forming conditions after 5 days. Significance determined by adjusted p-value of <0.05 with Benjamini-Hochberg correction. N=2-3 biological replicates. **(E)** HOMER annotation of Log2 Enrichment of KDM2A peaks in H2030-BrM3 tumor cell clusters segregated by their predicted activation or repression by KDM2A.

**Extended Data Figure 4:**
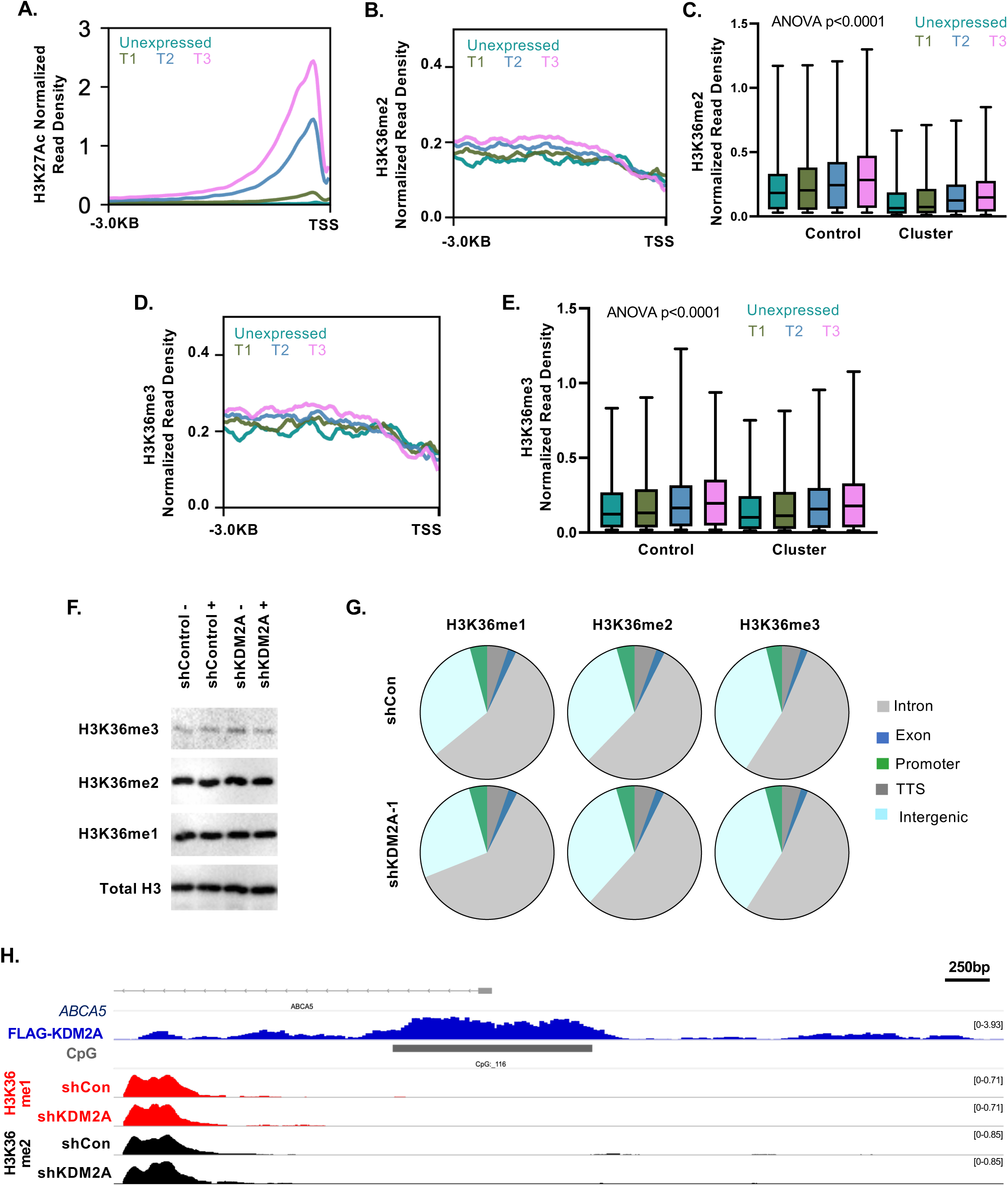
Chromatin remodeling by KDM2A in metastatic clusters. **(A)** Normalized read density (NRD) of H3K27Ac through the promoter region (−3kB to TSS) of genes in H2030-BrM3 cell clusters categorized by tertile of expression as in Figure 4B. N=4163 Unex., 5219 (T1), 5235 (T2), 5219 (T3) genes. Traces show average of 3 biological replicates taken at Day 5 after seeding in cluster forming conditions. **(B)** Profile plot of H3K36me2 CPM NRD as in Figure 4B. N=4163 Unex., 5219 (T1), 5235 (T2), 5219 (T3) genes. **(C)** Box-and-Whisker plots of NRD for traces underlying B and control conditions (data not shown). P-value by Kruskal-Wallis test with all comparisons p<0.0001 using Dunn’s multiple testing correction except: T0 cluster vs. T1 cluster (p=0.0025) and T0 control vs T1 control (p=0.5792). N=3642 (Unex.), 5397 (T1), 5398 (T2), and 5399 (T3) for control. **(D)** Profile plot of H3K36me3 NRD as in (B) N=4163 Unex., 5219 (T1), 5235 (T2), 5219 (T3) genes. **(E)** Box-and-Whisker plots of NRD for traces underlying D and control conditions (data not shown). P-value by Kruskal-Wallis test with all comparisons p<0.0001 using Dunn’s multiple testing correction except: T0 cluster vs. T1 cluster (p=0.0992), T0 control vs. T1 control (p=0.5396), T0 control vs. T2 cluster (p=0.9180), T1 control vs. T2 cluster (p>0.9999), and T2 control vs. T3 cluster (p>0.9999). N=3642 (Unex.), 5397 (T1), 5398 (T2), and 5399 (T3) for control. **(F)** Total protein levels of histone H3 and the indicated H3K36 modifications in shCon and shKDM2A H2030-BrM3 clusters +/− Dox for 5 days. **(G)** Proportion of peak summits annotated by HOMER to promoter, intergenic, intronic, exonic, or transcription termination site (TTS) regions in control or tumor cell cluster conditions. Proportions represent the average of peaks across N=4 biological replicates at Day 5 after. **(H)** IGV tracks of CPM NRD for the KDM2A suppressed gene *ABCA5*, with annotation of CGI, KDM2A chromatin binding peaks, and H3K36me1/2 peaks in shCon and shKDM2A-1 expressing H2030-BrM3 clusters. Traces are average of N=2 for FLAG-KDM2A and N=4 for H3K36 biological replicates.

**Extended Data Figure 5:**
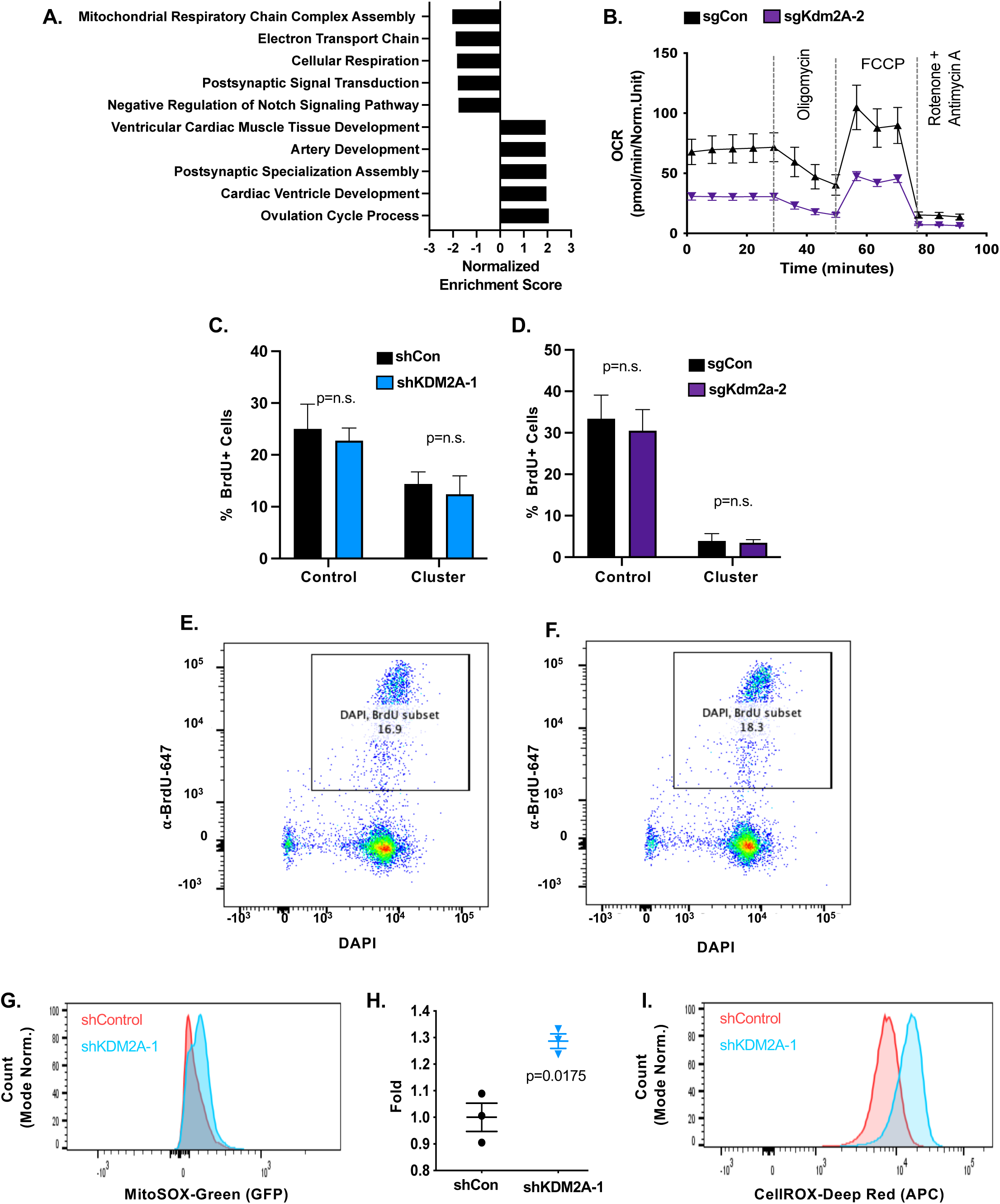
Cell biological effects of KDM2A inhibition in tumor cell clusters. **(A)** Top Gene Ontology - Biological Process pathway enrichments based on the differential expression of direct KDM2A targets following *KDM2A* knockdown in H2030-BrM3 clusters. All pathways shown have nominal p<0.05. **(B)** OCR was measured after 6 days of culturing LU-KP clusters with indicated sgRNAs N=6 biological replicates, each dot represents mean +/− SEM and normalized to cell number. **(C)** Quantification of BrdU positive H2030-BrM3 cells with the indicated shRNAs and grown under control or cluster forming conditions for 4 days. N=3, p>0.99 for both comparisons, using Mann-Whitney. **(D)** Quantification of BrdU positive LU-KP as in (C). N=3, p>0.99 using Mann-Whitney. **(E-F)** Representative flow cytometry gating from C. N= 8369 shCon cells in (E), N=8402 shKDM2A-1 cells in (F). **(G)** Intracellular mitochondrial superoxides (SOX) were measured using MitoSOX Green in H2030-BrM3 clusters with the indicated shRNAs after 4 days. Representative gating of MitoSOX, N=8603 shCon cells (red), N=8988 shKDM2A-1 cells (blue). Peaks are shown as normalized to mode. **(H)** MitoSOX data from H are normalized to the average value of shCon cells and p-value calculated by Welch’s t-test. (**I)** Representative gating of CellROX deep red dye from Figure 5L. N=25541 shCon cells (red), N=26385 shKDM2A-1 cells (blue). Peaks are shown as normalized to mode. Bar graphs show mean +/− SEM unless otherwise noted.

**Extended Data Figure 6:**
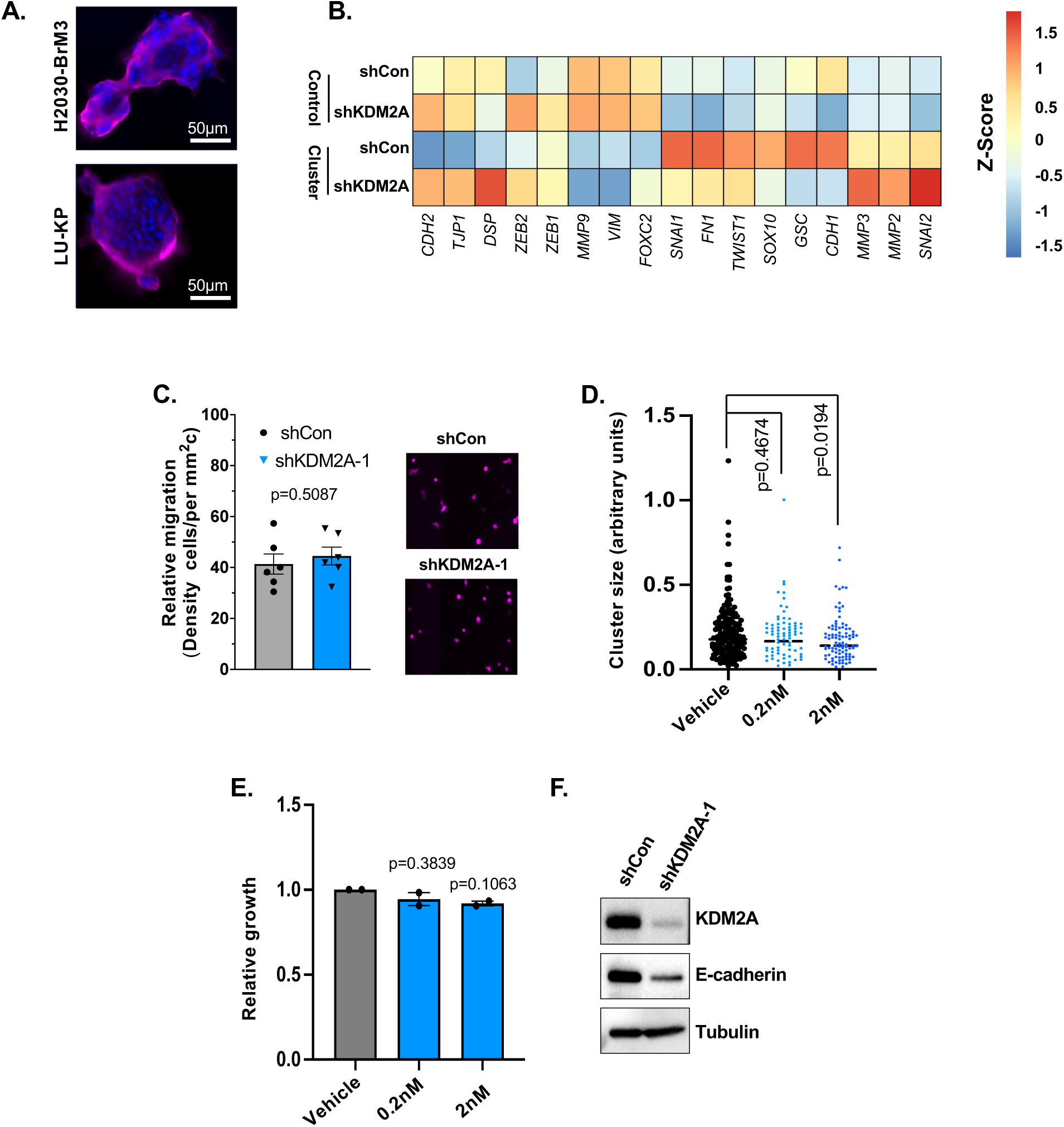
Effects of KDM2A knockdown on epithelial features in NSCLC cells. **(A)** Normalized expression of a curated list of genes involved in epithelial to mesenchymal plasticity and apical cell junctions. N=2-3 biological replicates. **(B)** Representative IF images of junctional permeability in H2030-BrM3 and LU-KP tumor cell clusters, based on Biotin labeling and AlexaFlour-647 streptavidin staining (pink). DAPI shown in blue. Scale=50μm. **(B)** Relative migration of H2030-BrM3 cells with the indicated shRNAs through a transwell barrier was measured after 24 hours (left). N=6 replicate images. P-value not significant by Mann-Whitney. Shown are representative images (right) for RFP positive cancer cells that have migrated across the barrier. **(D)** Cluster size measurements as in Figure 6C, of H2030-BrM3 cells treated with vehicle or ouabain octahydrate at the indicated concentrations for 6 days. P-value by Kruskal-Wallis test with Dunn’s multiple testing correction. N=2 biological replicates. **(E)** Relative growth for ouabain octahydrate treated clusters shown in (C) and measured as in Figure 1G. P-value by one-sided t-test against mean of vehicle. N=2 biological replicates. **(F)** Western blot of KDM2A and E-cadherin in H2030-BrM3 tumor clusters after Dox induction of shRNAs for 5 days. Alpha-tubulin shown as a loading control.

**Extended Data Figure 7:**
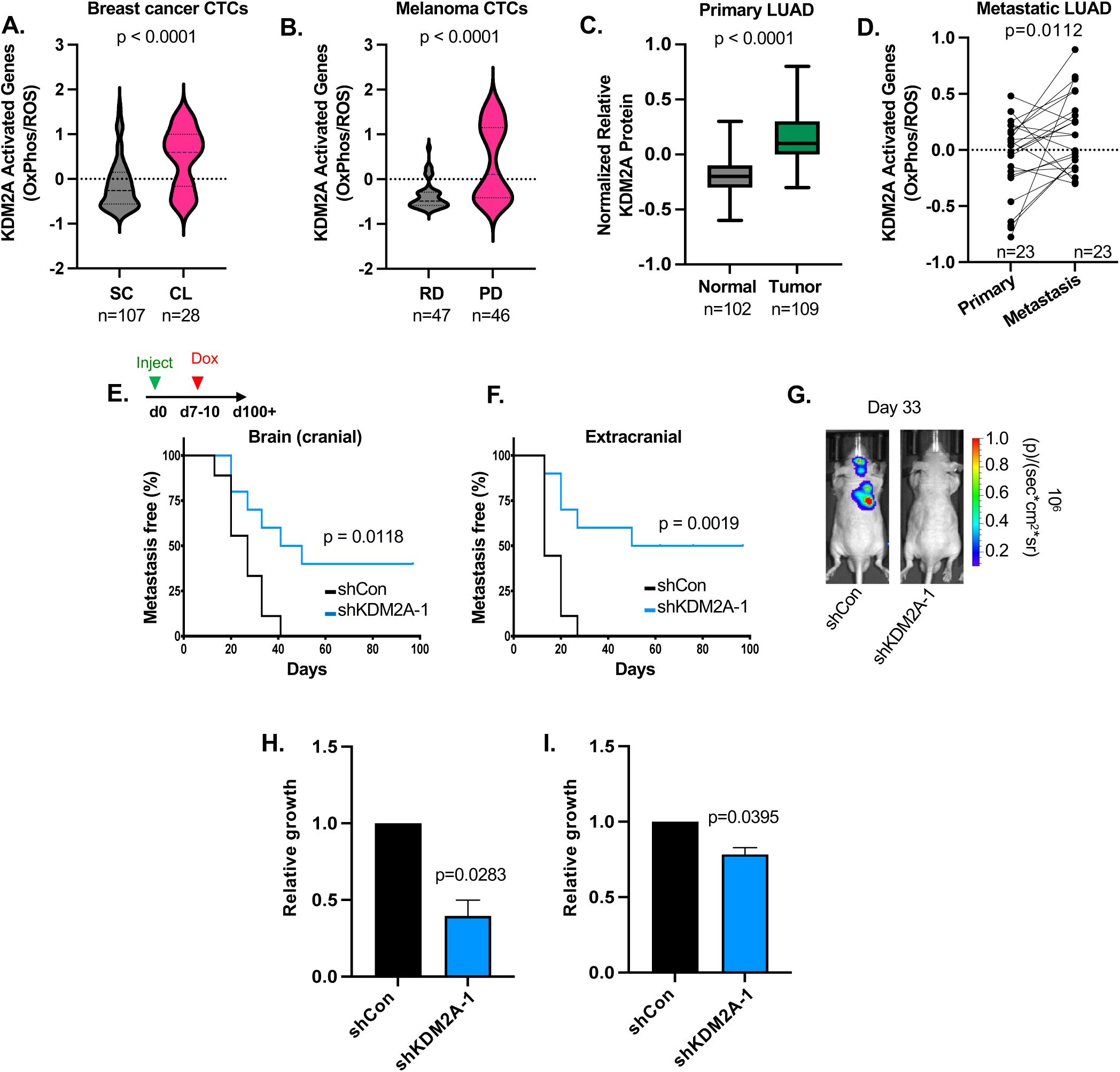
KDM2A mediates metastatic outgrowth. **(A)** Average expression of KDM2A bound and activated OXPHOS and ROS genes in individual CTCs from patients in Figure 7A, grouped by capture status as single cells (SC) or clusters (CL). N=107 single cells, 28 clustered cells. Data from GSE144495. P-value by Mann-Whitney. **(B)** Average expression of the KDM2A activated OXPHOS and ROS signature in human melanoma CTCs from patients with progressive metastatic disease (PD, N=46 cells) or treatment responsive disease (RD, N=47 cells). P-value by Mann-Whitney. Data from GSE157743. **(C)** Normalized relative protein levels of KDM2A from the Clinical Proteomic Tumor Analysis Consortium (CPATC) and Human Protein Atlas were plotted for primary lung adenocarcinomas vs. normal tissue. P-value by Mann-Whitney. Data from PDC000153. **(D)** Paired analysis of KDM2A OXPHOS and ROS signature expression in the primary and metastatic LUAD patient samples plotted in Figure 7B. P-value by Wilcoxen matched-pairs test. N=23 patients. Data from Zenodo record 10932811**. (E,F)** Mice on normal diet were injected with untreated H2030-BrM3 cells. 7-10 days after intracardiac injection, animals were switched to a Dox containing diet to induce the indicated shRNAs *in vivo*. Incidence of metastasis to the brain (E) or extracranial sites (F) was plotted as in Figure 7E and F. P-value by Mantel-Cox Log Rank Test. N= 9 shControl, 10 shKDM2A-1. **(G)** Representative BLI images on Day 33 of mice from (E) and (F). **(H)** Relative growth of PC9-BrM4 cells the indicated shRNAs was quantified based on luciferase activity after 12 days in culture forming conditions. P-value by one-sample t-test. N=3 per group. **(I)** Relative growth of MDA-BrM2 clusters with the indicated shRNAs was quantified as in (H). P-value by one-sample t-test. N=3 per group.

